# Human placental stem cells induce a novel multiple myeloid cell-driven immunosuppressive program that ameliorates proinflammatory CNS pathology

**DOI:** 10.1101/2025.06.14.659704

**Authors:** Heela Sarlus, Yuxi Guo, Stefan Bencina, Keying Zhu, Vijay Joshua Balasingh, Giulia Adriana Virgilio, Yufei Cheng, Jianing Lin, Jin-Hong Min, Irene Benito-Cuesta, Valerie Suerth, Paula Trigo Alonso, Olivia G Thomas, Urszula Rykaczewska, André Ortlieb Guerreiro Cacais, Maja Jagodic, Roberto Gramignoli, Robert A. Harris

## Abstract

Despite a growing interest in Amniotic Epithelial Cell (AEC)-based therapies, the immune responses triggered by AEC transplantation *in vivo* remain poorly characterized. In particular, how direct exposure to AECs within the central nervous system (CNS) shapes the local immune environment is currently unknown. Herein we describe a novel CNS-specific immunoregulatory pathway induced by intracisternal delivery of human AECs. Local immune responses induced by AECs in the brain led to recruitment of immunosuppressive Arginase 1^+^ (ARG1^+^) macrophages and a novel population of myeloid-derived suppressor cells with eosinophilic characteristics, which we term Eo-MDSCs. We further demonstrate that Eo-MDSCs produce Maresin 2 (MaR2), a specialized pro-resolving mediator (SPM) involved in the resolution of inflammation. In a mouse model of Multiple Sclerosis (MS), treatment of established disease with AECs induced immunological responses that resulted in reduced numbers of pathogenic macrophages and T helper (T_H_)17 cells, increased anti-inflammatory T cell subsets, and enhanced myelin phagocytosis, all of which led to functional recovery. These findings suggest that AEC therapy has the potential to target CNS-intrinsic inflammatory processes in MS, providing a strong rationale for translation into the clinic.

## Introduction

Chronic inflammation is a common denominator in many neurodegenerative diseases, which contributes to their pathogenesis. Multiple Sclerosis (MS) is an incurable autoimmune disease characterized by inflammation and demyelination, leading to progressive neurodegeneration. While T cells are central players in MS pathogenesis^1^, myeloid cells are crucial in the initiation and exacerbation of MS^2^. Myeloid cells in the brain predominantly comprise the central nervous system (CNS)-resident microglia and macrophages, as well as infiltrating monocyte-derived macrophages. During MS, myeloid cells infiltrate the CNS^3^ through meningeal routes^4,5^ where they contribute to neuronal damage by enhancing pathogenic T cell responses, secreting proinflammatory mediators^6^ and through active phagocytosis of axons^7^. Macrophages are prevalent in MS lesions^8–11^ and their numbers are traditionally known to correlate with demyelination and axonal damage^10,12–14^. However, over past decades it has become evident that myeloid cells have a remarkable capacity to resolve inflammation and to promote tissue repair in the brain. Consistent with this notion, we demonstrated that selectively inhibiting microglia proliferation^15^ or enhancing their myelin clearance capacity^16–18^ effectively induced disease remission in an experimental autoimmune encephalitis (EAE) model of MS. It is increasingly evident that current MS treatments targeting the adaptive immune arm exert their therapeutic effect indirectly by modulating myeloid cell activity^19,20^. Indeed, we and others have demonstrated that increased infiltration of immunosuppressive myeloid cells in the CNS reverts the pathogenic inflammatory processes leading to recovery in murine models of MS^21,22^. This multifaceted behaviour of myeloid cells attributed to their different activation states thus positions myeloid cells as important determinants of MS outcome.

AECs are stem cells derived from the innermost layer of the human placenta, offering multiple advantages over other cell types for cell therapy. These include their non-immunogenic and non-tumorigenic properties, their recovery through use of a non-invasive procedure with limited ethical concern, and their higher cell abundance. Since AECs do not express HLA class II antigens nor co-stimulatory molecules such as CD80 and CD86, acute rejection does not occur after their transplantation. Efficacious AEC-based therapies are reported across a wide range of disease models including hepatic disorders, Crohn’s disease and neurological conditions such as stroke and Parkinson’s disease^23–25^.

One emerging mechanism of action is their intrinsic immunomodulatory capacities. Although this has been partly investigated *in vitro*, whereby AECs suppress inflammatory responses in both innate and adaptive immune cells^26^, the immune cell dynamics triggered by AECs after their transplantation *in vivo* remain poorly understood. How direct AEC exposure shapes the immune environment within the CNS is a significant concept, given the ongoing clinical trials with AECs^24,25,27,28^, including treatment of neurological diseases that often requires direct delivery into the CNS. A detailed characterization of AEC-based immunomodulation in the CNS is therefore essential to understand their mechanisms of action and to ensure safe and effective clinical application. Herein we demonstrate that AEC delivery into the brain mobilizes the influx of immunosuppressive ARG1^+^ macrophages and a population of previously unrecognized myeloid-derived suppressor cells with eosinophilic characteristics, which we term Eo-MDSCs. By testing this premise in the mouse model of MS, we demonstrate that AEC-induced immunological response modulates T cell activity, promotes myelin phagocytosis and reduces spinal cord inflammation, ultimately leading to functional recovery.

## Results

### AEC suppress proinflammatory immune responses *in vitro* and *in vivo*

We first validated the immunomodulatory function of AECs on primary macrophages and microglia. The phenotype of AECs was confirmed by their expression of HLA-G, K8/18 (keratinocyte marker) and CD55 (**Figure S1A**). Proinflammatory cytokines were analysed in AEC-preconditioned microglia or control microglia challenged with lipopolysaccharide (LPS)/IFNγ for 24 hours. The qPCR analysis revealed that AEC-preconditioning reduced the expression of proinflammatory genes such as *Il6*, *Tnf*α and inducible nitric oxidase (*Inos*) in microglia (**Figure 1A**). The reduction of IL-6 and TNFα was verified at the protein level using a cytokine bead array (CBA) (**Figure 1B**). Notably, IL-10 levels were increased following AEC treatment (**Figure 1B**).

**Figure 1.**
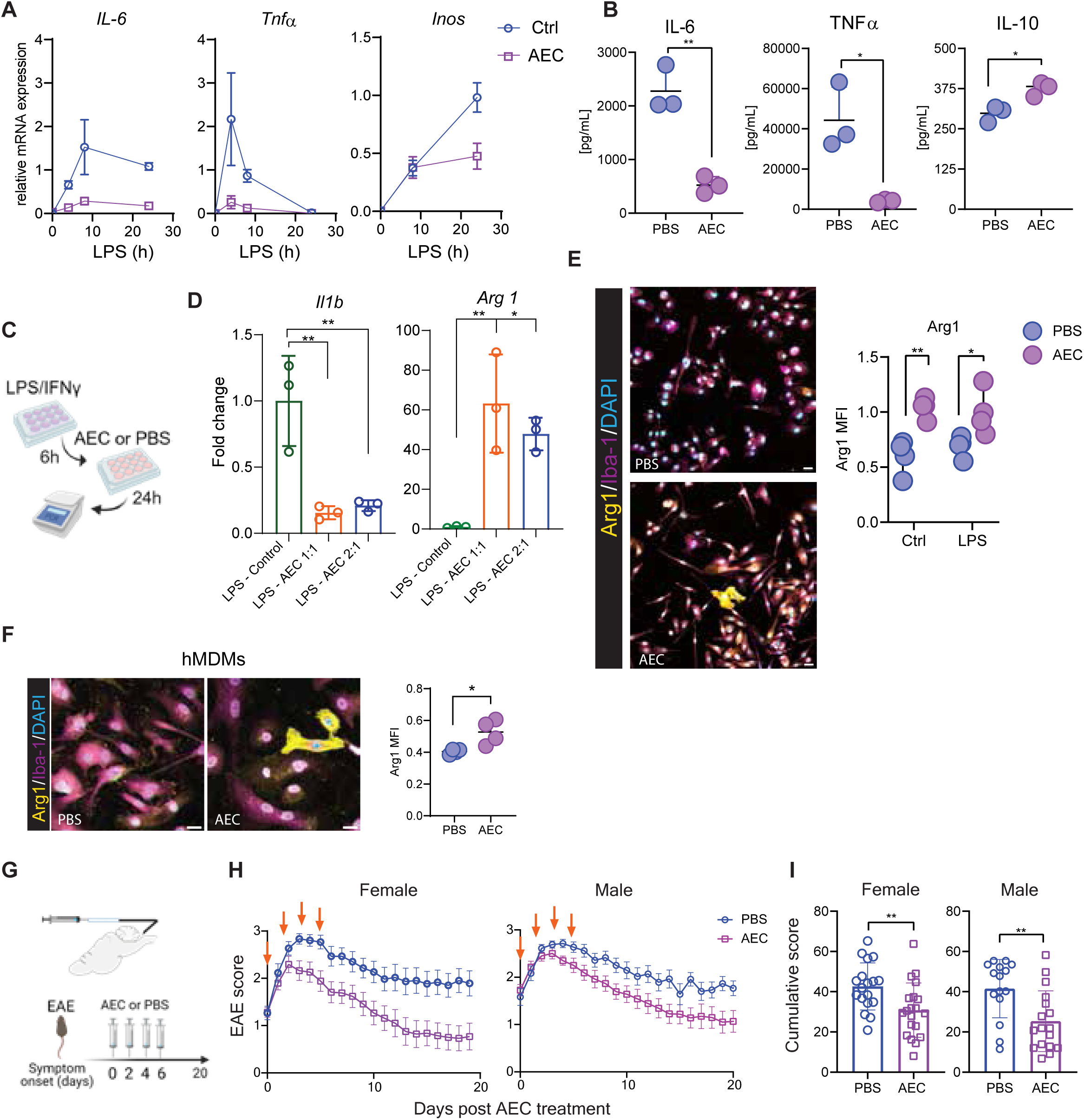
AEC reduces inflammatory macrophage responses *in vitro* and ameliorates EAE *in vivo*. **A** qPCR analysis of proinflammatory genes obtained from LPS/IFNγ stimulated microglia at different time points (4, 8, 24h) following preconditioning with or without AECs. Data are presented as mean ± SD; statistical analysis: unpaired Student’s t-test. **B** Secreted cytokines in the conditioned media obtained from cell culture after 24 hours of LPS/IFNγ stimulation. Data are presented as mean ± SD; statistical analysis: unpaired Student’s t-test. **C** Schematic illustration of with LPS stimulation followed by AEC preconditioning. **D** qPCR of LPS-stimulated macrophages exposed to PBS (control) or AEC at 1:1 or 2:1 macrophage:AEC ratios. Data are presented as mean ± SD; statistical analysis: one-way ANOVA followed by Tukey’s multiple comparison test. **E** Arg1 and Iba-1 immunofluorescence of macrophages preconditioned with or without AEC. Quantification was performed using Image J using default B&W thresholding. Data are presented as mean ± SD; statistical analysis: two-way ANOVA followed by Tukey’s multiple comparison test. **F** Representative expression of Arg1 in human monocyte-derived macrophages (hMDMs) and its quantification in the right panel. **G** Schematic illustration of AEC or PBS treatment paradigm at EAE symptom debut. **H** EAE disease score in female and male mice. Data are presented as mean ± SEM. The figure presents data from two independent experiments in each female and male mice. **I** EAE cumulative score for female and male mice. Data are presented as mean ± SEM; statistical analysis: Mann Whitney test. *p<0.05, **p<0.01, ***p<0.001

We next examined the capacity of AECs to reduce inflammatory responses in LPS-stimulated macrophages (**Figure 1C**). As expected, AEC treatment decreased the gene expression of the proinflammatory cytokine IL-1β, while the expression of Arginase 1 (*Arg1*), a marker of immunosuppressive macrophages, was markedly increased at gene (**Figure 1D**) and protein levels (**Figure 1E**). The increase in ARG1 was confirmed in human monocyte-derived macrophages (hMDMs) following AEC treatment, suggesting a conserved immunosuppressive response by macrophages to AEC (**Figure 1F**). Based on these results, we hypothesized that intracisternal (i.c.) delivery of AECs would lead to clinical recovery in a mouse model of neuroinflammation.

We first delivered PHK206-labelled AECs i.c. and detected them in the brain borders, including the ventricles, both at 3h and 3 days post injection (p.i.) (**Figure S1B**), confirming their retention in the brain. Subsequently, we examined the therapeutic impact of AEC on established neurological MS-like disease. When each mouse had developed an EAE disease score of ≥1 they received AEC injections every other day, 4 injections in total (**Figure 1G**). AEC therapy significantly reduced the macroscopic signs of clinical disease in both female and male mice, as indicated by reduced cumulative EAE scores (**Figure 1H and 1I**). Taken together these data demonstrate that AECs are efficient in reducing inflammation *in vitro* and therapeutically ameliorating symptoms of already established MS-like disease *in vivo*.

### Intracisternal AEC delivery induces CNS influx of immunosuppressive myeloid cells

We next examined the immunosuppressive immune responses elicited by AECs in the brain. We first analysed myeloid phenotypes in the brain at EAE peak (**Figure 2A**). AEC treatment substantially increased the numbers of myeloid cells (**Figure 2B-2D**, gating strategy **Figure S1C**) that consisted of monocytes, macrophages and a population of highly granular (HG) cells which were unique to AEC-treated brains. This immunological response was local in the brain and absent in the spleen, blood, bone marrow (BM) and skull BM at the time of EAE peak (**Figure S1D, S2**). HG myeloid cells characterized by the signature side-scatter (SSC)^high^F4/80^lo^MHCII^lo^ expressed Ly6C but not Ly6G, which phenotypically excludes them as being neutrophils (**Figure 2C, Figure S1C**). Using Giemsa staining we observed that most HG cells had ring-shaped nuclei, with some containing cytoplasmic granules (**Figure 2E**). We performed scRNAseq on sorted HG cells to define their transcriptional signature. We analysed 2858 HG cells sorted from the brain as SSC^high^CD45^+^ (PBS-treated: 917 cells; average genes 2226, AEC-treated: 1941 cells; average genes 1344) and identified five distinct HG clusters (**Figure 2F and 2G**).

**Figure 2.**
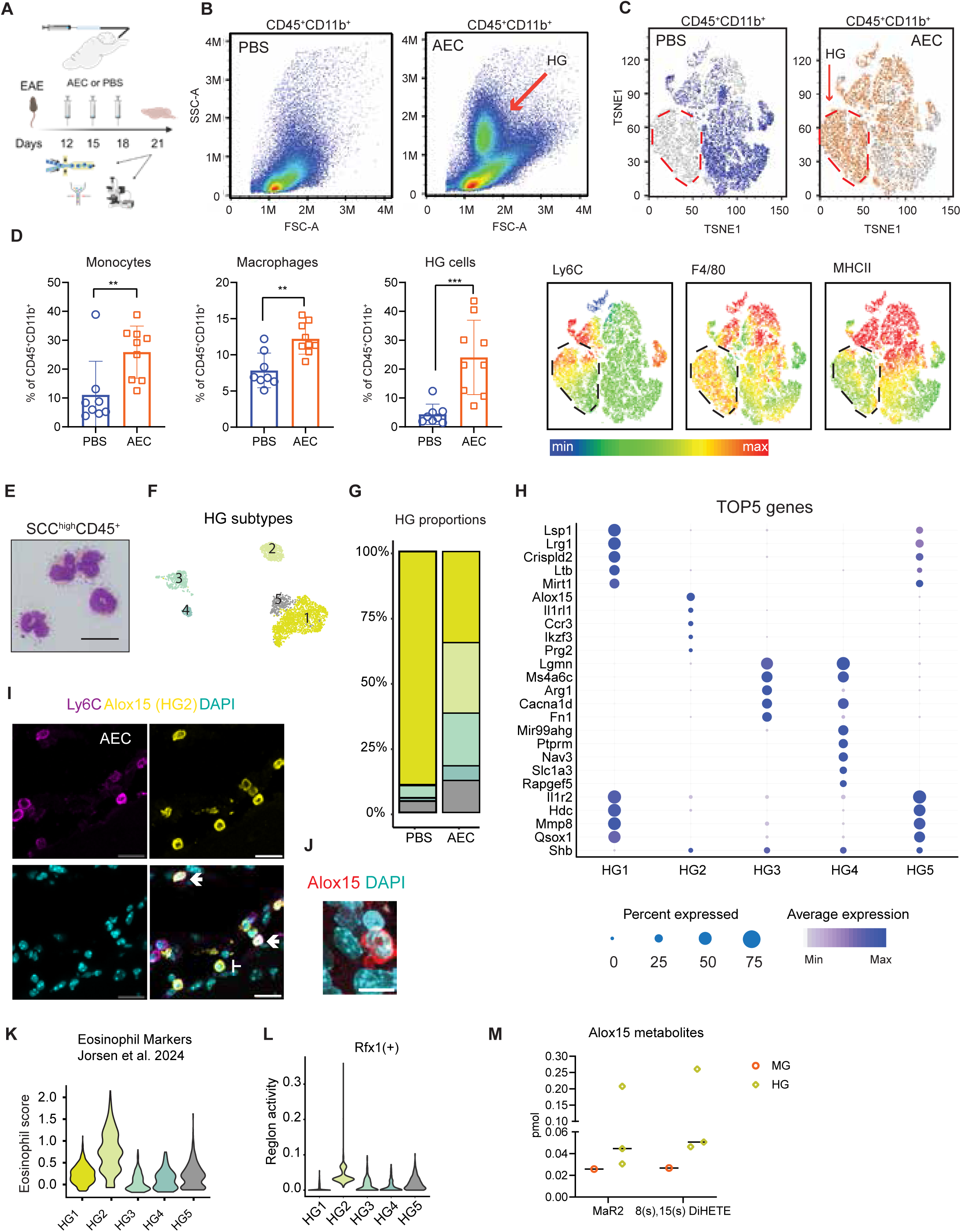
Intracisternal AEC delivery induces CNS influx of immunosuppressive myeloid cells. **A** Schematic depicting intracisternal delivery of AEC or PBS every other day, three times starting on day 12. Brains were collected three days post last AEC challenge. **B** FACS plot gated on myeloid cells in AEC- or PBS-treated EAE brains. **C** tSNE and feature plots generated in Flowjo of myeloid cells using the default settings, indicating the expression of different myeloid markers in AEC- or PBS-treated. **D** Quantification of myeloid subsets in the brain. n=8 (PBS), n=9 (AEC). Data are presented as mean ± SD; statistical analysis: unpaired Student’s t-test. **E** Giemsa staining of sorted SSC^high^CD45^+^ high granular cells. Scale bar=50µm. **F** UMAP projection of 2858 SSC^high^CD45^+^ high granular cells sorted from mouse brain three days after last AEC challenge and **G** their proportions. Cells are colored according to identified clusters. **H** Dot plot of top5 hallmarks. **I** Representative ALOX15 immunofluorescence indicating Ly6C expression. Scale bar=40µm. **J** Representative ALOX15 immunofluorescence in AEC-treated brains indicating HG2 consists of cells with ring-shaped nuclei. Scale bar=20µm. **K** Comparison of HG clusters with previously published eosinophil markers. **L** Violin plot illustrating transcription factor Rfx1 regulon activity in HG clusters. **M** ALOX15 metabolites in HG and microglia subsets. n=3 per condition. *p<0.05, **p<0.01, ***p<0.001.

HG1 and HG5 consisted of neutrophils (**Figure S3A**), and HG5 had reduced expression of genes such as *Lsp1*, *Lrg1* (**Figure 2H**), which are linked to neutrophil chemotaxis and differentiation^29,30^. Overall, neutrophil proportions were not affected by AEC treatment (**Figure S1E**). HG3 belonged to the monocyte/macrophage lineage and expressed *Ly6c2*, *Cd14*, *Csf1r*, and *Adgre1* (**Table 1**) with high expression of *Arg1* (**Figure 2H**). HG4 was a small cluster reminiscent of microglia, expressing genes such as *Tmem119*, *P2ry12*, *Sall1* and *Hexb* (**Table 1**). We focused on HG2, which was exclusive to AEC-treated brains (**Figure 2G**). Using arachidonate 15-lipoxygenase (ALOX15) immunostaining, a top gene expressed in HG2 (**Figure 2H**), we identified that HG2 consisted of cells with ring-shaped nuclei with variable Ly6C expression (**Figure 2I and 2J**), in accordance with the FACS data. Since HG2 have similar morphology and low RNA content (**Figure S3B**) as immature neutrophils^31^ and eosinophils^32^, we compared the transcriptome of HG2 to the genomic signature of the respective immune cells^33^. Our analysis revealed that the HG2 transcriptome had more overlap with eosinophils (**Figure 2K, Figure S3C**) than with neutrophils (**Figure S3A**). Furthermore, Single Cell Regulatory Network Inference and Clustering (SCENIC) analysis identified Rfx1, a transcription factor involved in eosinophil development^34^, to have higher regulon activity in HG2 (**Figure 2L**). Immature myeloid cells are also MDSCs, so we therefore compared the HG2 transcriptome with previously published polymorphonuclear (PMN) and monocytic (M) MDSC signatures. Our analysis revealed little overlap (**Figure S3D and S3E**)^35,36^, suggesting that HG2 are immature myeloid cells or MDSCs with eosinophilic features, which have not been described previously.

Given the high expression of ALOX15 in HG, we assessed ALOX15 metabolites in sorted HG cells using lipidomics, using microglia as controls. Interestingly, levels of Maresin 2 (MaR2), a specialized pro-resolving lipid mediator (SPM) with proven anti-inflammatory function, and 8(s)-15(s)-DiHETE, a chemoattractant for eosinophils, were higher in HG cells compared to controls (**Figure 2M**).

Altogether, we demonstrate that intracisternal delivery of AECs increases immunosuppressive myeloid cells in the brain characterized by *Arg1*^+^ monocytes and a novel anti-inflammatory MDSC population, Eo-MDSCs.

### AEC treatment transiently recruits myeloid cells into the CNS

We next performed immunostaining to localize ARG1^+^ macrophages and Eo-MDSCs in the brain at peak of EAE disease. ARG1^+^ macrophages were only noted sporadically, if at all, in PBS-treated brains. However, in AEC-treated brains the numbers of ARG1^+^ macrophages were remarkably increased, mainly localizing to brain barriers including the meninges, ventricles, and in subependymal areas (**Figure 3A**, **Figure S3F**) and choroid plexi (**Figure 3B**), hereafter collectively referred to as ‘brain borders’. Morphologically, ARG1^+^ macrophages were distinct from ARG1^-^ counterparts (**Figure S3G**). ALOX15^+^ Eo-MDSCs were also identified within the brain borders (**Figure 3A, Figure S3H**). The accumulation of these immunosuppressive myeloid cells in the meninges may have a disease-modifying impact as meningeal inflammation correlates with relapses and a severe disease course^37,38^.

**Figure 3.**
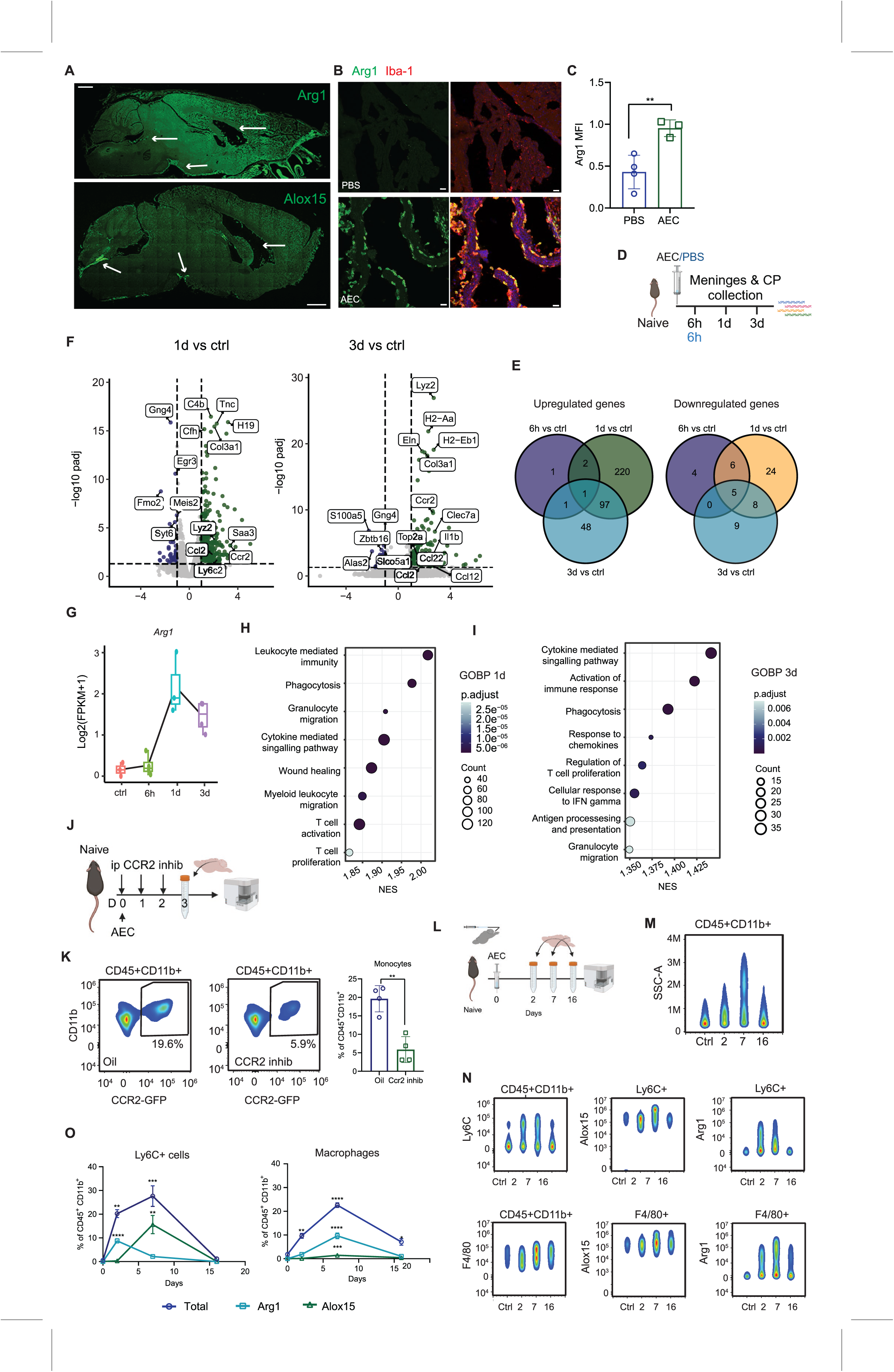
AEC treatment transiently recruits myeloid cells into the CNS. **A** Representative image of ARG1 and ALOX15 immunofluorescence in the brain. Scale bar=1000µm. **B** Representative expression of ARG1 in choroid plexus and **C** its quantification. Data are presented as mean ± SD; statistical analysis: unpaired Student’s t-test. Scale bar= 40µm. **D** Experimental design of brains for bulkRNAseq **E** Venn diagram indicating upregulated and downregulated genes 6h, 1day (1d) or 3 days (3d) post-AEC challenge. Control mice received PBS. n=4 per group. **F** Volcano plot for 1d and 3d post- AEC challenge. **G** Box plot indicating *Arg1* expression at different time points post-AEC treatment. **H-I** GOBP terms for the same time points. **J** Experimental design CCR2 inhibition experiment. **K** FACS plot indicating CCR2^GFP^ monocytes in AEC-treated mice with or without CCR2 inhibition. Data are presented as mean±SD; statistical analysis: unpaired Student’s t-test. n=4 per condition. **L** Experimental design of myeloid cell kinetics following AEC treatment. **M-O** FACS plot indicating myeloid subsets after different days post-AEC injection: **M** SSC gated on myeloid cells (CD45^+^CD11b^+^), **N** Upper panel: Ly6C-expressing myeloid cells indicating ALOX15, and ARG1 subsets within Ly6C^+^ population. Lower panel: F4/80-expressing myeloid cells indicating ALOX15, and ARG1 subsets within F4/80^+^ population. **O** Quantification of different ARG1^+^ and ALOX15^+^ myeloid subsets at 2, 7 and 16 days post-AEC administration. Data is presented as mean±SD; statistical analysis: One-way ANOVA with Bonferroni’s multiple comparison. Ctrl n=4; 2d n=4; 7d n=6; 16d n=5

We next asked whether the influx of these myeloid cells in the CNS is driven by the release of cytokines and chemokines in brain borders following AEC administration. To this end, we isolated the leptomeninges and choroid plexus for bulk RNAseq at 6h (immediate response), 1 or 3 days p.i. (**Figure 3D**). After 6h, only five genes were significantly upregulated, whereas this number was 320 and 147 genes after 1 and 3 days p.i., respectively (**Figure 3E**). The number of downregulated genes was low at all time points (**Figure 3E**). As expected, AEC treatment increased the expression of monocyte transcripts *Lyz2, Ccr2,* and *Ly6c2* and their attracting chemokines *Ccl2, Ccl12, Ccl22* (**Figure 3F**). Interestingly, *Arg1* expression was already elevated by day 1 p.i. and persisted through to day 3 (**Figure 3G**). The Gene Ontology (GO) analysis indicated enrichment related to myeloid migration (**Figure 3H, 3I**). Pharmacological inhibition of CCR2 significantly attenuated AEC-induced influx of monocytes, demonstrating CCR2- driven recruitment of monocytes into the brain (**Figure 3J, 3K**).

Having established that AEC treatment drives the influx of immunosuppressive myeloid cells, we then investigated how long these cells remain in the brain. We analysed brain myeloid responses at 2, 7 or 16 days post-AEC injection using flow cytometry (**Figure 3L**). Control naïve mice received PBS instead of AECs. We observed an influx of monocytes peaking at 2 days p.i., and a subset of these monocytes (Ly6C^+^) expressed ARG1^+^ (**Figure 3M, 3N, 3O**). By day 7, the numbers of ARG1^+^ monocytes declined, but instead the numbers of ARG1^+^ macrophages (F4/80^+^) peaked, indicating a monocyte-to-macrophage maturation (**Figure 3M, 3N, 3O**). The kinetics of ALOX15^+^ Eo-MDSCs were different. At day 2 p.i. highly granular cells were absent, but they peaked at day 7 (**Figure 3M**), with a concomitant increase in ALOX15^+^ Eo-MDSCs (**Figure 3M, 3N**). By day 16, the numbers of both ALOX15^+^ Eo-MDSCs and ARG1^+^ macrophages declined to basal levels (**Figure 3O**). We confirmed this in EAE brains at the end of the observation period, with the numbers of ARG1^+^ macrophages being comparable between AEC and PBS groups (**Figure S3I**) and in naïve mice that received 3 AEC injections (**Figure S3J, S3K, S3L**).

Taken together, our data indicate that AEC therapy induces a transient accumulation of ARG1^+^ macrophages and ALOX15^+^ Eo-MDSCs in the brain, creating a temporally restricted immunosuppressive milieu, particularly at the brain borders, capable of modulating neuroinflammatory processes.

### Intracisternally delivered AECs reprogram monocytes towards an anti-inflammatory phenotype

We next investigated how the influx of AEC-induced myeloid cells impacts neuroinflammation during EAE. We profiled CD45^+^ cells from the brains of AEC- or PBS-treated mice at the time of EAE peak using scRNAseq and excluded high granular cells (**Figure 4A**). After quality control, 30677 cells (PBS: 15906 cells; average genes 2129, AEC: 14777 cells; average genes 2191) were analysed. We identified both innate and adaptive immune cells in the brain (**Figure S4A-S4C**), obtaining 6 microglial clusters, but their proportions were not altered by AEC treatment at this time point (**Figure S4D and S4E**). Intriguingly, monocytes and macrophages were major responders to AEC treatment. We identified nine clusters and denoted them as six monocyte clusters and three macrophage clusters based on the presence or absence of *Ccr2* and *Ly6c2* expression (**Figure 4B-4D**).

**Figure 4.**
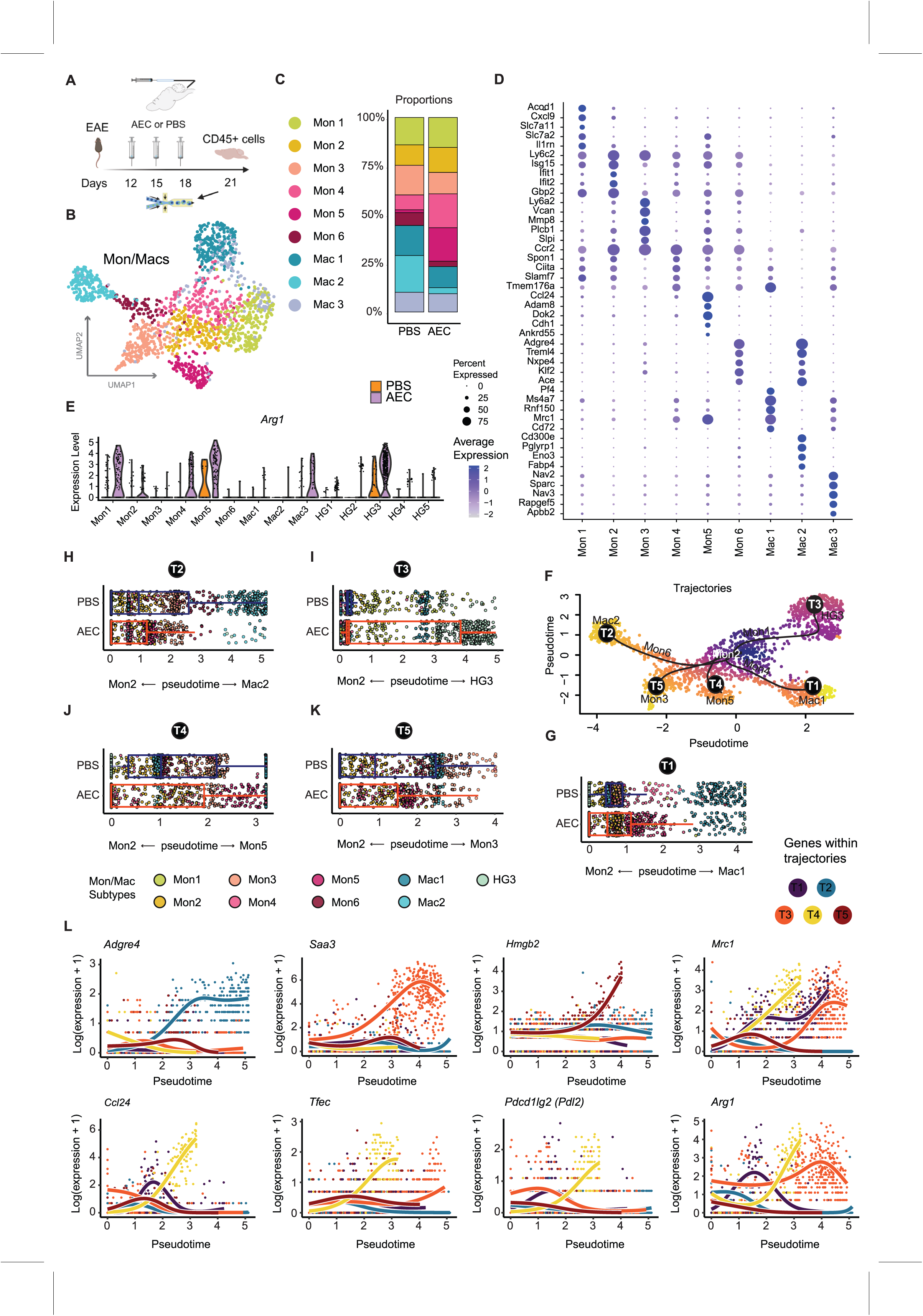
AEC treatment increases Arg1^+^ macrophages in the brain. **A** Schematic depicting the experimental design. **B** UMAP of 1560 macrophages/monocytes cells and **C** their proportions in PBS- or AEC-treated brains. **D** Top5 genes in each Mon/Mac cluster. **E** Violin plot indicating DEG *Arg1* expression in different monocyte/macrophage subsets. **F** Phenotypic trajectory inferred by Slingshot indicating five potential trajectories starting with Mon2. **G-K** Distribution of monocyte/macrophage cells in pseudotime across the five main transcriptional trajectories depicting **G** Mac1, **H** Mac2, **I** HG3, **J** Mon5, and **K** Mon3. **L** Expression profile of selected DEGs across pseudotime axis.

The clusters that increased substantially in response to AEC treatment included Mon4 and Mon5. Cells in Mon5 expressed high levels of *Mrc1, Ccl24* and *Arg1* (**Figure 4D-4E**), which are induced after stimulation with cytokines such as IL-4 and IL-10 to drive immunosuppressive macrophage polarization^39^. Conversely, the clusters that were substantially reduced in response to AEC treatment were Mon6 and Mac2. Two of the top 5 Mac2 genes included *Pglyrp1* and *Fabp4* (**Figure 4D**), which have been ascribed pathogenic effects in models of inflammatory diseases. Pglyrp1 deficiency in myeloid cells was protective against the development of EAE^40^. Similarly, inhibition of FABP4, an adipokine secreted by proinflammatory synovial macrophages, alleviated the development of rheumatoid arthritis^41^. In MS patients, serum FABP4 levels correlate with disease severity^42^, altogether indicating a proinflammatory role of Mac2 cells.

We next performed trajectory analysis using Slingshot to dissect how monocytes phenotypically transition to different states depending on treatment. HG3, which was closely associated with monocyte clusters in the combined HG and non-HG datasets (**Figure S4A**), was also included in this analysis. We discerned five distinct trajectories starting with Mon2 (**Figure 4F**). In the PBS-treated group, most monocytes phenotypically transitioned to either Mac2 or Mon3 (**Figure 4H and 4K**), with proinflammatory *Adgre4*^43^ and *Hmbg2* (**Figure 4L**) being upregulated genes in the respective trajectories. In contrast, monocytes in the AEC-treated group predominantly transitioned to Mon5 or HG3 (**Figure 4I and 4J**). In the trajectory leading to Mon5 (T4), anti-inflammatory genes such as *Arg1* and *Ccl24*, as well as transcription factor *Tfec* which is inducible by IL-4 in macrophages^44^, were increased (**Figure 4L**). Furthermore, *Pdl2,* an immune checkpoint that restricts effector T cell responses was also upregulated in the Mon5 trajectory (**Figure 4L**). In the HG3 trajectory (**Figure 4I**), one of the highly upregulated genes was serum amyloid A (*Saa3*). Although SSAs have been implicated in many inflammatory conditions^45^, their function in polarizing immunosuppressive macrophages, characterized by elevated *Arg1* and *Mrc1*^46^ expression, supports our findings (**Figure 4L**).

Collectively, these data pinpoint that AEC treatment reduces the transition towards proinflammatory macrophages, while increasing monocyte transitioning towards *Arg1*^+^ macrophages (Mon5, HG3), which have immunosuppressive functions and are known to promote recovery in EAE^47^.

### AEC treatment induces Tregs and T_H_2 cells in the brain

Immunosuppressive myeloid cells are known to modulate T cell responses and to thereby influence MS-like disease course^22^. We thus evaluated T cell responses at the EAE peak in the brain (**Figure 5A**). We consistently observed a reduction in the proportion of T_H_17 cells in AEC-treated brains (**Figure 5B**, gating strategy in **Figure S5A**), whereas those of anti-inflammatory Tregs and T_H_2 increased (**Figure 5B-5C**). IL-4 is a cytokine mainly produced by T_H_2 cells and its deficiency aggravates EAE^48,49^, and conversely the adoptive transfer of IL-4-overexpressing T cells ameliorates EAE^50^. This indicates that T_H_2 responses in EAE have a protective role. Intriguingly, cellular interaction analysis using CellChat revealed that in AEC-treated brains, T_H_2 cells communicated with Mon5 through IL-4 (**Figure 5D**). This interaction is required for the persistence of *Arg1* macrophages^51^. Given that ARG1^+^ monocytes emerge early in response to AEC therapy, we hypothesized that they promote IL-4^+^ T_H_2 polarization. To test this, we injected activated T cells derived from MOG-immunized 2D2 mice i.c. into PBS- or AEC-treated naïve recipients (**Figure 5E**). T cells were injected on day 6 post-AEC injection, a timepoint when ARG1^+^ macrophages/monocytes are present in the CNS, and the brains were analysed 4 days later. As expected, polarization to IL-4^+^ T_H_2 cells was increased in brains preconditioned with AECs (**Figure 5F, Figure S4F**), highlighting a potential mechanism whereby ARG1^+^ macrophages drive anti-inflammatory T cell responses and contribute to neuroprotection.

**Figure 5.**
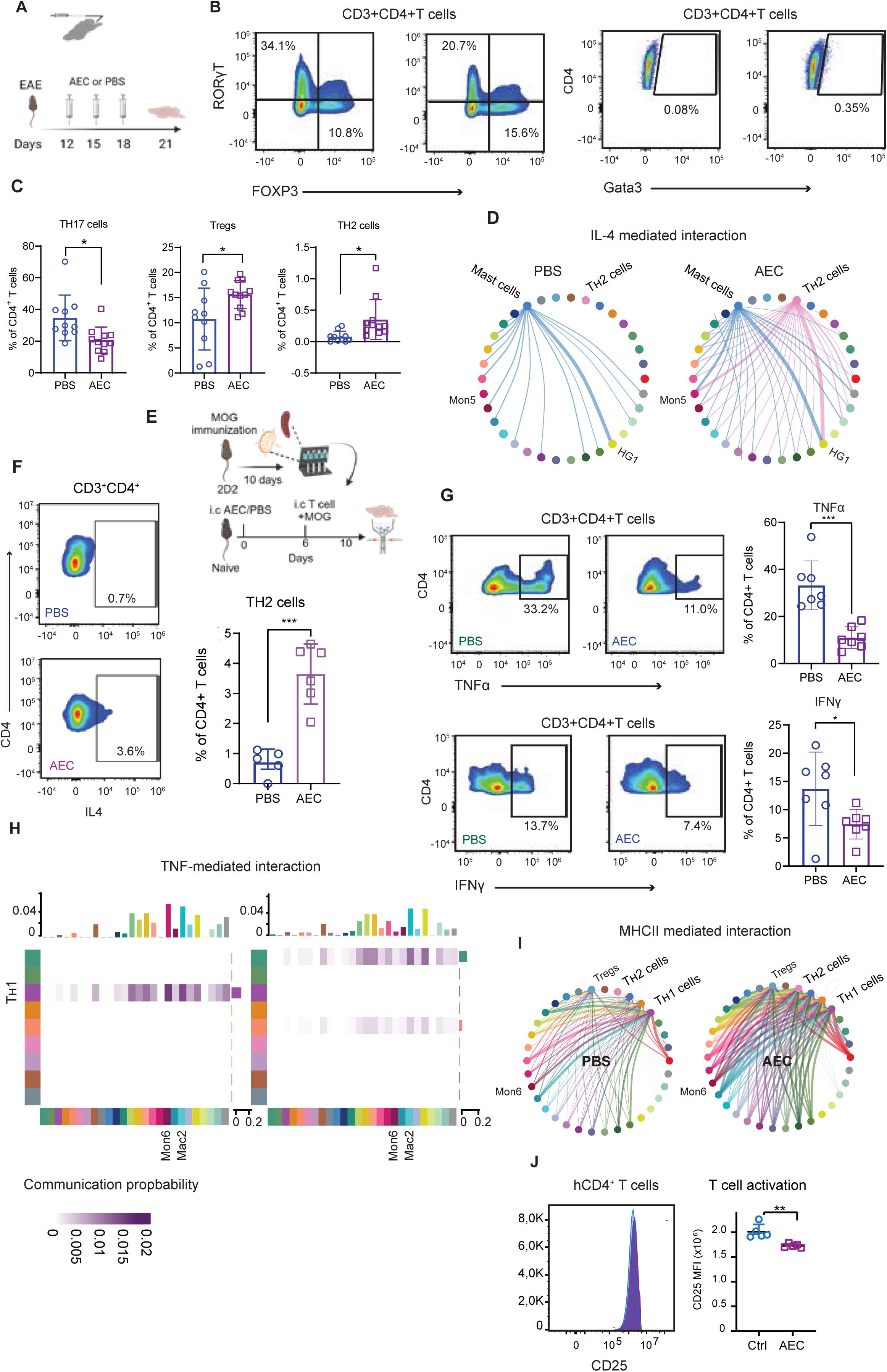
AEC treatment impact on brain T cells. **A** Schematic depicting experimental design. **B** FACS plots indicating RORγt, FOXP3, and GATA3 expressing CD4^+^ T cell subset and **C** their quantification, with averages indicated in the respective FACS plot. n=10 (PBS), n=11 (AEC). Data are presented as mean ± SD; statistical analysis: unpaired Student’s t-test. **D** Interaction strength inferred by CellChat analysis. CellChat interaction plot indicating IL-4- mediated interaction. **E** Experimental design for T cell transfer experiment. **F** FACS plot gated on CD4^+^ T cell subsets indicating IL-4 expression, with quantifications in the right panel. Data are presented as mean ± SD; n=5 PBS and n=6 AEC. statistical analysis: unpaired Student’s t-test. **G** FACS plots indicating TNFα and IFNγ cytokine production in CD4^+^ T cells and their quantification to the right panel, with averages indicated in the respective FACS plot. Data are presented as mean ± SD; statistical analysis: unpaired Student’s t-test. **H** Interaction plot inferred by CellChat indicating TNF-mediated interaction with T_H_1, Mon6 and Mac2 subsets highlighted. **I** MHCII-mediated interaction between myeloid and T cell subsets inferred by CellChat analysis. **J** Histogram indicating CD25 expression in CD4^+^ T cells derived and expanded from MS patients. n=5 (Ctrl), n=5 (AEC). Data are presented as mean ± SD; statistical analysis: unpaired Student’s t-test. \**p* < 0.05; \*\**p* < 0.01; \*\*\**p* < 0.001

We also evaluated the levels of the proinflammatory cytokines TNFα and IFNγ, which are known to exacerbate EAE^52,53^, at the peak of EAE using flow cytometry. The proportions of IFNγ- and TNFα-producing CD4^+^ T cells were reduced in AEC-treated brains (**Figure 5G**). These findings were verified *ex vivo* in MOG-restimulated T cells obtained from lymph nodes of MOG-immunized 2D2 mice, which express MOG-specific T cell receptors (**Figure S5E-S5F**). It has been reported that T cell-derived TNFα exacerbates EAE by recruiting monocytes into the CNS^52^. Indeed, CellChat analysis revealed that in PBS-treated brains TNF-mediated interactions were strongest between the sender T_H_1 cells and Mon6 and Mac2 receivers (**Figure 5H**). Intriguingly, these interactions were strongly reduced in AEC-treated brains (**Figure 5H**), with a concomitant reduction in the above-mentioned myeloid subsets (**Figure 4C**), thus corroborating the role of T cell-derived TNFα in inflammatory monocyte recruitment^52^.

CellChat analysis further revealed that MHC class II-mediated interaction between T and myeloid cells was remarkably enhanced in AEC-treated groups. Notably, in PBS conditions Tregs received minimal myeloid input, whereas in AEC conditions this interaction was significantly enhanced (**Figure 5I**).

Collectively, these data indicate that following AEC treatment, not only proinflammatory T cell responses are reduced, but also myeloid cell interaction with anti-inflammatory T cells is enhanced, thereby contributing to the suppression of neuroinflammation in EAE.

We next studied T cell responses to AEC-preconditioned hMDMs. We obtained T cells derived from MS patients and after clonal expansion we co-cultured CD4^+^ memory T cells with AEC- or control-preconditioned hMDMs under stimulation with αCD28/αCD3. Interestingly, the expression of CD25, a late activation marker for memory CD4^+^ T cells, was decreased in T cells co-cultured with AEC-preconditioned hMDMs (**Figure 5J**).

Altogether, our data demonstrate that AEC treatment reduces pathogenic T cell responses and increases anti-inflammatory T cell subsets, which are associated with their therapeutic effect.

### AEC treatment attenuates proinflammatory myeloid cells and spinal cord pathology

We next studied immune responses in the spinal cord following AEC treatment at EAE peak using flow cytometry (**Figure 6A**). Intriguingly, the myeloid response in the spinal cord was distinct from that in the brain. Overall, the proportion of CX3CR1^+^F4/80^+^macrophages was reduced in AEC-treated spinal cords (**Figure 6B**, gating strategy in **Figure S6A**). However, a higher proportion of these macrophages expressed CD206^+^, a marker expressed by immunosuppressive macrophages (**Figure 6C**). We also identified HG cells in AEC-treated spinal cords. In addition, we observed a reduction in microglia/macrophage infiltrates in spinal cord lesions in AEC-treated animals (**Figure 6F**), while homeostatic microglia marker P2RY12 expression was higher (**Figure 6G**), altogether indicating reduced spinal cord inflammation following AEC-treatment.

**Figure 6.**
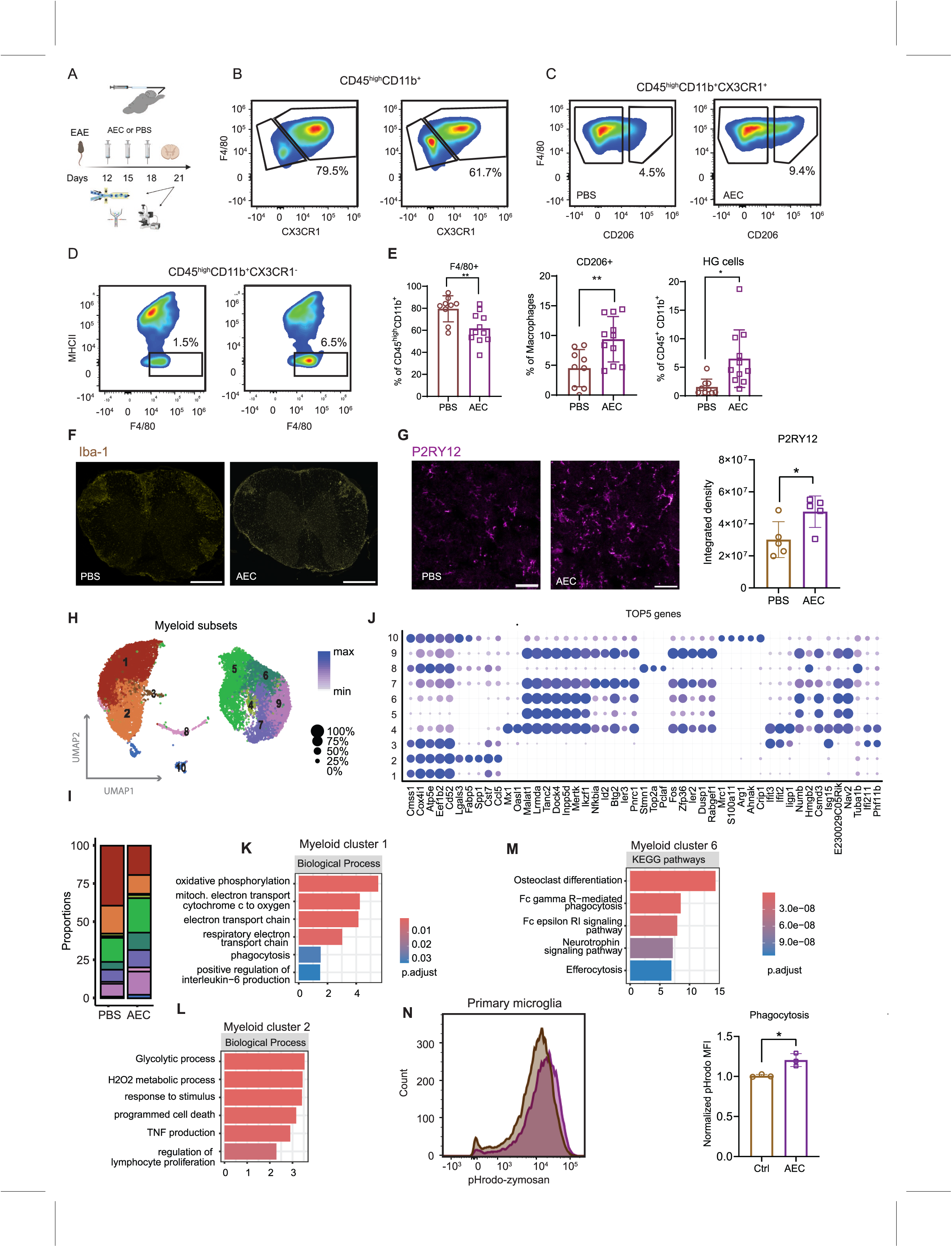
AEC treatment attenuates proinflammatory myeloid cells and spinal cord pathology. **A** Experimental design. **B** FACS plots indicating CX3CR1^+^ F4/80^+^ macrophages within CD45^high^ myeloid subset in the spinal cord. **C** FACS plots indicating CD206^+^ subset within macrophages in the spinal cord. **D** FACS plots indicating F4/80^lo^MHCII^-^ subset high granular (HG) cells in the spinal cord. **E** Quantification of each population with averages indicated in the respective FACS plot. Data is presented as mean ± SD; n=9 PBS and n=11 AEC. Statistical analysis: unpaired Student’s t-test. **F** Iba-1 immunofluorescence in the spinal cord of PBS- and AEC-treated EAE spinal cords. Scale bar= 500 µm. **G** P2RY12 immunofluorescence in the spinal cord of PBS- and AEC-treated EAE spinal cords, with quantification to the right. Data are presented as mean±SD; statistical analysis: unpaired Student’s t-test. n=5 PBS and n=5 AEC. Scale bar= 40 µm. **H** UMAP of 13228 myeloid cells and **I** their proportions in PBS- or AEC-treated spinal cords. **J** Top5 genes in each myeloid cluster. **K** GOBP term Myeloid cluster 1. **L** GOBP term for Myeloid cluster 2. **M** KEGG pathways for Myeloid cluster 6. **N** FACS plot indicating zymosan fluorescence intensity (uptake) in primary microglia treated with AEC-condition media or media (Ctrl). Data is presented as mean ± SD; statistical analysis: unpaired Student’s t-test. n=3 \**p* < 0.05; \*\**p* < 0.01; \*\*\**p* < 0.001

To further characterize myeloid responses we performed scRNAseq of spinal cord-derived CD45^+^ cells at EAE peak (**Figure 6A**). After quality control, 14105 cells (PBS: 6009 cells; average genes 1680, AEC: 8096 cells; average genes 1934) were analysed, yielding 10 myeloid clusters (**Figure 6H-6J**). We identified several clusters (1, 2, 5, 6, 7) of disease-associated microglia (DAM) (**Figure S6B**), interferon responsive microglia (cluster 3, 4), proliferating microglia (cluster 8), immediate response microglia (cluster 9) and cluster 10, which resembled brain Mon5 expressing *Mrc1* (CD206) and *Arg1* (**Figure 6J**). Interestingly, the proportions of clusters 1 and 2 were considerably reduced in AEC-treated spinal cords. Both clusters consisted of DAM-expressing mitochondrial and glycolysis genes which were previously demonstrated to drive neuroinflammation through increased production of reactive oxygen species^54^. Indeed, the GO analysis of these respective clusters confirmed enrichment related to mitochondrial electron transport chain and glycolytic processes (**Figure 6K-6L**). Furthermore, KEGG pathway (**Figure S6C**) enrichment terms included neurodegenerative diseases, neurodegeneration and the HIF-1 signalling pathway, altogether indicating their pathogenic role in EAE^54^. Conversely, cluster 6 was highly increased in AEC-treated spinal cords. This cluster had low expression of MHC II genes (**Figure S6D**) and KEGG analysis indicated enrichment in pathways related to neurotrophin signalling, phagocytosis, and efferocytosis (**Figure 6M**), which are key steps in resolution of inflammation.

We next tested the phagocytic function of microglia by exposing primary microglia treated with AEC-conditioned media to pHrodo-labelled zymosan particles. Intriguingly, zymosan uptake was increased in microglia preconditioned with AEC-conditioned media (**Figure 6N**). To further explore whether AEC-induced macrophages could effectively clear myelin debris, a key function mediated by immunosuppressive macrophages to facilitate remyelination and tissue repair^55,56^, we exposed myeloid cells isolated from PBS- or AEC- treated brains at the EAE peak to pHrodo-labelled myelin debris for 60 min *ex vivo* (**Figure S6E**). Overall, a higher proportion of myeloid cells phagocytosed myelin debris in AEC-treated brains, and macrophages constituted the majority of these cells (**Figure S6F**). HG cells were not involved in phagocytosis (**Figure S6G**). Consistent with brain, a higher proportion of macrophages engulfed myelin debris in the AEC-treated spinal cords (**Figure S6H**).

## Discussion

In this study we uncover a novel CNS-localized immunoregulatory circuit induced by AECs which enable mobilization of immunosuppressive myeloid cells into the brain through the choroid plexus and meninges. These myeloid cells comprise both ARG1^+^ immunosuppressive macrophages and anti-inflammatory Eo-MDSCs that decrease pathogenic macrophage and T cell activities and produce regenerative factors, ultimately leading to immunosuppression and functional recovery (**Figure 7**). While the *prophylactic* effects of AEC treatment have previously been reported in EAE^57,58^, we significantly extend these studies and demonstrate that myeloid cells are the primary responders to i.c. treatment with AECs.

**Figure 7.**
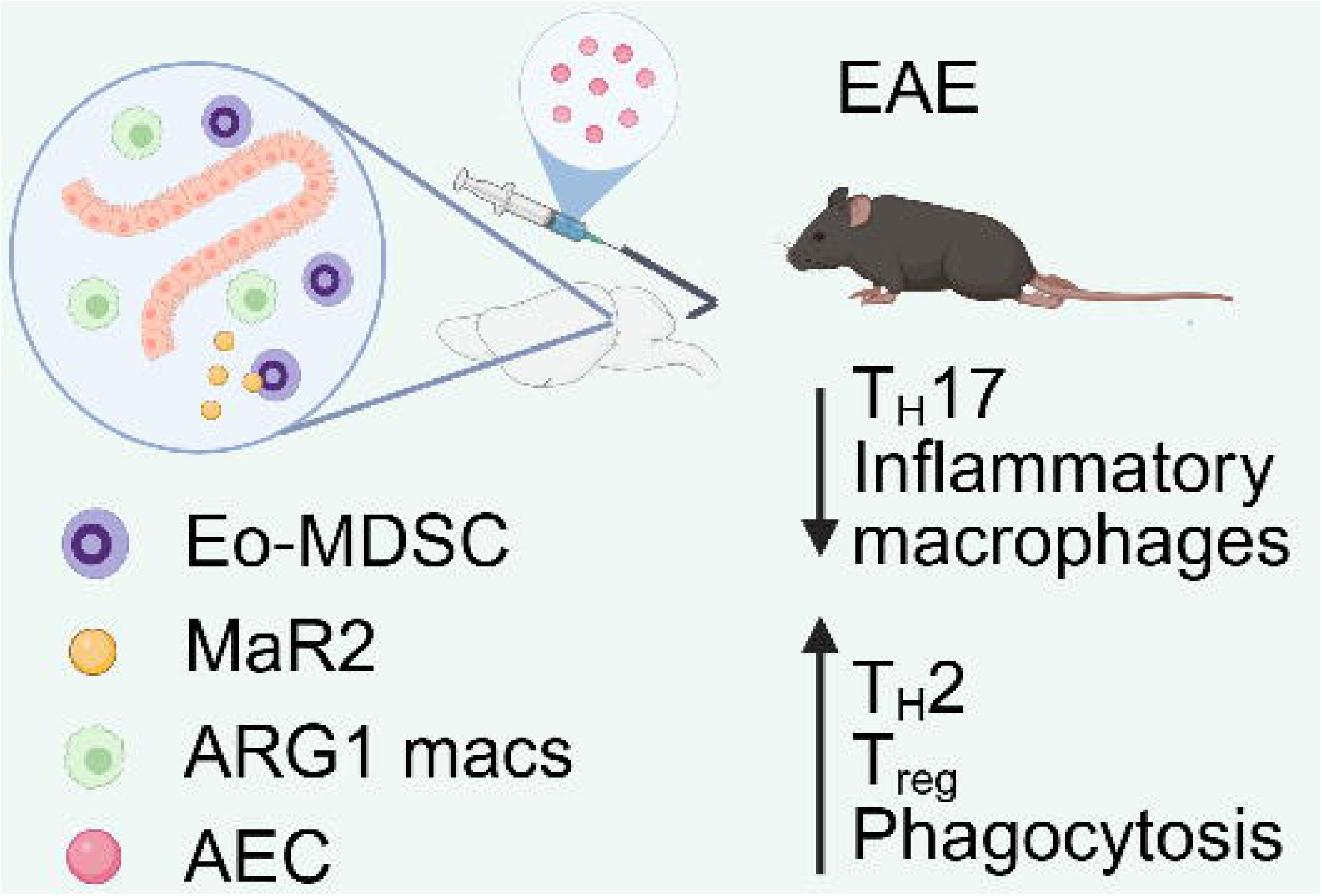
**Summary of the work**. Intracisternal delivery of human placenta-derived AECs induces a CNS-specific immunoregulatory circuit characterized by accumulation of immunosuppressive macrophages ARG1^+^macrophages and a novel population of myeloid derived suppressor cells with eosinophilic characteristics termed as Eo-MDSCs. These cells secrete maresin 2 (MaR2), a specialized pro-resolving mediator involved in resolution of inflammation. In EAE mice with established diseases, this AEC-induced immunological response modulates T cell responses by increasing polarization towards T_H_2 and T_regs_ while decreasing the pathogenic T_H_17 subsets. In addition, pathogenic microglia subsets are reduced as well as AEC-recruited macrophages promote myelin debris clearance important for tissue repair.

In neuroinflammatory diseases such as MS, myeloid cells are implicated in causing injury to the CNS but also in promoting tissue repair^22,59^, a dichotomous response governed by different myeloid activation states. Our study demonstrates that AEC therapy, by modulating myeloid cell function in the CNS, has the potential to target multiple pathological processes in MS. Firstly, broad rim lesions enriched with iNOS^+^ macrophages correlate with progression in MS^60,61^. Since ARG1 and iNOS compete for the same substrate, AEC-induced ARG1^+^ macrophages have the capacity to directly counteract the proinflammatory effects of iNOS^+^ macrophages, thereby targeting pathogenic processes. Secondly, ARG1^+^ macrophages are known to increase in numbers during the resolution phase of EAE to promote tissue repair^47,62^, and enhanced phagocytosis is one underlying mechanism. Our data suggest that through enhanced myelin debris clearance, a critical step required for efficient remyelination^63^, AEC-induced immunosuppressive macrophages have the potential to promote remyelination processes in the CNS. Finally, these immunosuppressive macrophages can have protective function by counteracting the excessive MS inflammation driven by T_H_1 and T_H_17 cells. We observed a significant reduction of pathogenic T cell responses. T cell-derived TNFα promotes the activation and infiltration of inflammatory monocytes/macrophages and autoreactive T cells in the CNS^52,53^. However, TNFα produced by other cells can have beneficial functions as blocking TNFα leads to detrimental outcomes in both MS patients and in EAE^53^. That AEC treatment specifically attenuated T cell-derived TNFα and modified their interactions with myeloid cells suggest that AEC therapy effectively targets pathogenic MS mechanisms. Furthermore, locally produced IL-4 in the brain effectively ameliorates EAE^50^, probably by driving immunosuppressive macrophage polarization.

Induction of Eo-MDSCs in the brain is an interesting feature of AEC therapy. We demonstrate that these cells produce MaR2, a SPM which has not been previously studied in EAE, but which promotes the resolution of inflammation in other settings. However, MaR1, which is similar to MaR2, reduces neuroinflammation and increases numbers of Tregs in EAE^64^, suggesting overlapping mechanisms. Due to the lack of specific markers and their shared features with other myeloid cells, the specific role of MDSCs in MS may have been overlooked. PMN-MDSC and M-MDSCs have conflicting roles described in MS^65,66^, although their protective functions in restricting pathogenic T_H_1 and T_H_17 responses and reducing tissue damage are evident from many studies^65,67^. The ambiguity of MDSC function in MS might thus reflect their heterogeneity, where the functional state of MDSCs and the stage of the disease in which they appear may dictate the outcome. In a therapeutic context, glatiramer acetate, which has been used for treating relapsing-remitting MS, induces M-MDSCs^68^. However, Eo-MDSCs have never been reported before in any MS context. Further studies are therefore needed to elucidate their specific function in both EAE and MS.

Brain barriers are important entry sites for proinflammatory myeloid cells and pathogenic T cells to access the brain and spinal cord during MS^5,69,70^. An association between meningeal inflammation and chronic MS is evident in several studies, often indicating the presence of higher numbers of lymphocytes and T cell infiltrates in the meninges, which correlate with cortical demyelination and a severe disease course. That meningeal inflammation occurs early in MS^71^ and its model^5,70^ and correlates with the ensuing damage suggests its involvement in MS pathogenesis. By modulating immune activity in the CNS gateways such as the choroid plexus and the meninges, AEC-induced anti-inflammatory myeloid cells have a strong potential to reduce CNS-intrinsic inflammation and to promote regenerative processes.

A limitation of the current study is that we did not track the transplanted AECs in the CNS. Although few AECs labelled with a commercially available dye were detected in the ventricles 3 days after AEC injection, the dye was not stable, and the cells could not be clearly located. The fate of AECs in the CNS has yet to be unequivocally determined by tagging them using more stable tracking methods.

In summary, our study greatly advances our understanding of local CNS responses to stem cells in the brain, providing valuable insights into the underlying mechanisms of their therapeutic effects. We demonstrate that AEC therapy not only orchestrates myeloid activation into immunoregulatory subtypes, mitigating excessive inflammation driven by T cells and macrophages, but also promotes functional recovery and activation of regenerative processes. This provides a rationale for clinical translation of AEC therapy into several neurodegenerative disease settings.

## Materials and Methods

### Experimental animals

Male and female C57BL/6NTac mice (Taconic) were bred at the Comparative Medicine Department at Karolinska University Hospital, Sweden. Animals were maintained in a pathogen-free and climate-controlled environment with regulated 12 h light/dark cycles. All mice used for experiments were adults between 2-4 months of age and had access to chow and water *ad libitum*. The experiments were performed in accordance with the Swedish National Board of Laboratory Animals and the European Community Council Directive (86/609/EEC) and the local ethics committee of Stockholm North under the ethical permit 61327-22.

### Induction of Experimental Autoimmune Encephalomyelitis (EAE)

For EAE induction, mice were immunized subcutaneously in the dorsal tail base with 40µg MOG emulsified in complete Freund’s adjuvant (Chondrex, 7027) containing 200µg *Mycobacterium tuberculosis* per mouse, under isoflurane (Baxter, 1001936040) anesthesia. MOG emulsion was prepared using the POWER-Kit according to the manufacturer’s instructions (BTP emulsion, Malmö, Sweden, https://btbemulsions.com). In addition, mice received i.p injections of 200ng pertussis toxin (Sigma, P7208) at the time of EAE induction and 48h post-immunization. Mice were weighed and fed with wet food daily from Day 7 post-immunization, and scored according to the following scheme: 0=no clinical score, 0.5=reduced tail tension, 1=drooping tail (no tonus), 1.5=drooping tail with clumsy gait, 2=hindlimb paraparesis, 2.5=one hindlimb dragging with paraparesis in the other hindlimb, 3=hindlimb paralysis, 3.5=hindlimb paralysis with forelimb paraparesis, 4=tetraplegia or moribund, 5=death. Mice were euthanized if they exhibited >25% weight loss or prolonged tetraplegia for more than 3 days without signs of recovery. The clinical symptoms were scored daily in a blinded manner.

### AEC isolation

Human AECs were isolated from freshly delivered full-term placentae as previously described^72^. Briefly, the amnion membrane was peeled off from the underlying chorion layer and pre-weighed pieces of amnion membrane were digested with TrypLE 10X (cat.# A12177-01, Life Technologies) for 30 min at 37C° in a rotator. The membranes were washed with Plasmalyte solution (cat. # AFE0324D, Baxter) to further release the cells. The cell suspension was collected and centrifuged at 300g for 10 min at 4°C and filtered through a 100 µm cell strainer. Cell number and viability were obtained through the Trypan Blue Exclusion method (cat.# 15250061, Gibco). Ten million AECs were resuspended in 1 mL of CryoStor10 (cat.# c2874, Merck) and stored in liquid nitrogen until required.

### Intracisternal injection of AECs

Cryopreserved AECs were washed with PBS and centrifuged at 350g for 5 min at 4°C. The cells were resuspended in PBS to obtain 5x10^6^ cells/mL. For intracisternal (i.c) injections mice were anesthetized under isoflurane with the head bending forwards to expose the aperture of the cisterna magna. A 27G dental needle (Terumo, DN-2721) with the tip (around 3.5mm) bent at an angle of approximately 40° was connected to a Hamilton syringe via a polyethylene tube and inserted into the cisterna magna and 10μL of solution was slowly injected during approximately 10s.

For treatment studies, after reaching a score of ≥1 each mouse received approximately 6x10^5^AECs suspended in PBS every other day 3-4 times. Control mice underwent the same procedure but only received PBS. For tracking AECs, the cells were labelled with PHK26 dye (mini26-1KT, Sigma) according to the manufacturer’s instructions and subsequently injected i.c as described above.

### Isolation of myeloid cells from brain/spinal cords

Brains or spinal cords from C57BL/6NTac mice were dissected after cardiac perfusion with cold PBS under overdose of isoflurane anaesthesia. The brain tissue was first mechanically dissociated and subsequently digested in 5mL papain (Worthington, LS003126; 1:100 diluted in L15 medium) supplemented with DNase I (Roche, 10104159001, 0.2mg/mL). The brain homogenate was then incubated at 37 in a water bath for 20 min, with pipetting in between every 10min. After terminating the enzymatic reaction by adding 20mL cold HBSS, the cell homogenate was passed through a 40µm cell strainer and centrifuged at 350g for 5 min at 4 . The cell pellets were subsequently resuspended in 20mL 37% isotonic Percoll (Sigma-Aldrich, P1644) in HBSS and centrifuged at 800g (acceleration 4, deceleration 0) for 10min at 4°C. Subsequently, the myelin was removed (used as myelin debri), the cells were washed with PBS and pelleted at 350g for 5 min at 4°C for downstream analysis.

### Primary cell culture

To generate BMDMs, femurs obtained from C57BL/6NTac were flushed and the resulting single-cell suspensions were washed with PBS and centrifuged at 350g for 5 min at 4°C. The pellet was resuspended in macrophage medium containing Dulbecco’s modified Eagle’s medium (DMEM; Sigma-Aldrich, D6046) supplemented with 10% heat-inactivated fetal bovine serum (FBS; Sigma-Aldrich, F7524), 10 ng/mL recombinant mouse M-CSF (R&D Systems, 416-ML), 2mM l-glutamine (Sigma-Aldrich, G7513), 100U/mL penicillin, and 100μg/mL streptomycin (Sigma-Aldrich, P4458). The cells were cultured in a T175-cell culture flask (Sarstedt, 83.3912.502) and half the cell medium was replaced on day 4 and fully changed on day 6. The cells were harvested after 7-8 days using trypsin/EDTA solution (Gibco, 25300096), plated for 24h and subsequently used in experiments.

For primary microglia the mixed glial cell pellets (obtained above) were resuspended in microglia medium consisting of DMEM/F12 medium, 10% FBS, 20ng/mL recombinant mouse M-CSF (R&D Systems, 416-ML), 2mM l-glutamine (Sigma-Aldrich, G7513), 100U/mL penicillin, and 100μg/mL streptomycin (Sigma-Aldrich, P4458). Mixed glial cells were cultured and expanded for approximately 14 days with medium changes twice a week. When the cells were confluent, they were digested with trypsin as above. Pure microglial cultures were obtained using magnetic beads with anti-CD11b MicroBeads (Mitenyi Biotec, 130-049-601) according to the manufacturer’s protocol.

Cryopreserved AECs were thawed in DMEM low glucose medium (cat#11054020, Gibco) supplemented with 5% Stemulate® human platelet lysate (Biolife Solutions). After centrifugation at 300g for 5 min at 4°C, cell pellets were reconstituted in 1X HBSS and counted with Trypan Blue Exclusion method. The pellets were resuspended in DMEM low glucose medium supplemented with 5% Stemulate® human platelet lysate, 1% MEM Non-essential Amino Acid Solution 100X (cat# 11140050, Gibco) and 1% L-Glutamine 220mM (cat# 25030081, Gibco). Cells were cultured in collagen precoated T75-cell culture flask (Sarstedt, 83.3911.002) for two weeks, with medium change every 3 days. Subsequently, the cells were harvested using trypsin/EDTA solution (Gibco, 25300096) as above. AECs were resuspended in DMEM/F12 medium, 10% FBS, 2mM l-glutamine (Sigma-Aldrich, G7513), 100U/mL penicillin, and 100μg/mL streptomycin (Sigma-Aldrich, P4458). AECs were seeded at the densities of 400, 000 AECs per well in a 24- well plate (Nunc 142475, Thermofisher Scientific) and 20,000 cells per well in a 96-well plate (Cat. # 3603, Life Sciences) for 24h and used in subsequent experiments.

### Cell stimulation and treatment

To induce an inflammatory response in macrophages and microglia, we applied 100ng/mL LPS and 20ng/mL IFNγ. The cells were pre-incubated with AEC at 1:2 AEC:macrophage/microglia ratio for 3 or 5 days in either macrophage or microglia medium (see above). For qPCR and cytokine analyses the cells were plated in 12-well plates at a density of 3x10^5^ cells per well. The cells were harvested at different time points post-LPS challenge.

### Human monocyte-derived macrophages

Blood from healthy donors was obtained through Karolinska University Hospital. Procedures were conducted under ethical permits approved by the Swedish ethical review authority. Peripheral blood mononuclear cells (PBMCs) were isolated from blood, positively selected for CD14 and differentiated into MDM. Briefly, blood mixed with PBS was carefully added to Ficoll-Paque PLUS (Cytiva). After centrifugation, mononuclear cell layers were collected and washed in PBS. PBMCs were resuspended in 90% FBS and 10% DMSO and stored at −80°C. For positive selection of monocytes, MACS was performed using anti-human-CD14 MicroBeads (Miltenyi Biotec). Isolated monocytes were cultured in MDM medium composed of RPMI 1640 medium GlutaMAX (Thermo Fisher Scientific), 10%FBS, 50ng/ml M-CSF(PeptroTech), 10ng/ml GM-CSF(PeproTech), 0.055mM 2-mercaptoethanol (Gibco), 1% Penicillin/Streptomycin with half the medium changed every 48 hours until day 12. MDMs were cultured with or without AECs for 5 days in MDM medium and thereafter used for immunostaining (see Immunofluorescent staining) or co-cultured with T cells (see below).

### Human T cell preparation

Natalizumab-treated persons with MS had been diagnosed relapsing-remitting MS according to the McDonald 2010 criteria. Patients were recruited at the Karolinska University Hospital, Stockholm, Sweden and the Academic Specialist Center, Stockholm, Sweden. All human subjects gave written informed consent to blood sampling and the study was approved by the Swedish Ethical Review Authority’s Stockholm Ethics Board (no. 04-252/1-4 and no. 2009/2107-3112).

Briefly, CD45RA-negative cells were isolated using CD45RA microbeads (Miltenyi) from cryopreserved natalizumab-treated MS patient PBMC according to the manufacturer’s instructions and stained with CSFE (Thermofisher Scientific). After stimulation with antigen beads at a ratio of 10:1, CD45RA-negative cells were incubated for 8 days at 37°C in 5% CO_2_. Subsequently, cells were stained with Live/Dead Aqua viability dye (Thermofisher Scientific), CD3 APC-Cy7 and CD4 PerCP-Cy5.5 antibodies (BioLegend) and the LiveCD3^+^CD4^+^CSFE^DIM^ population was sorted using a SH800S Cell Sorter (Sony). For expansion of T cells, single LiveCD3^+^CD4^+^CSFE^DIM^ T cells were sorted into wells of 96-well U-bottom plates containing 2x10^5^ irradiated allogeneic PBMC (45 Grays) per well in T cell media (IMDM 5% human serum (Merck), 100U/ml penicillin, 100μg/ml streptomycin, 2mM L-glutamine) containing Phytohaemagglutinin (Thermofisher Scientific) and 20U/ml human IL-2 (Peprotech) and expanded for 2 weeks. T cells were thereafter co-cultured with MDMs preconditioned with AECs (1:3 AEC: T cell ratio) for 3 days under stimulation with CD28/CD3.

### Flow cytometry

Single cell suspensions from brain or cells *in vitro* (above) were stained with live dead Live/Dead Fixable Near-IR or Yellow Dead Cell Stain Kit (Invitrogen, L34976 or L34959, 1:500) in the presence of Fc Block (Invitrogen, 14-0161-83, 1:200). For the analysis of CNS immune cells, the following antibodies were used with incubation time of 30 min at 4 C°: CD45-BUV496 (clone 30-F11, 749889, BD Biosciences), CD11b-Percp-Cy5.5 or CD11b BV650 (clone M1/70, 101228, Biolegend), Ly6C-AF700 or Ly6C-PE (clone HK1.4, 128024, Biolegend), Ly6G-AF647 (clone 1A8, 127609, Biolegend), F4/80-BV421 (clone BM8, 123137, Biolegend), MHCII-Spark-UV387 (clone m5/114.15.2, 107669, Biolegend), Clec7α-Percp-Eflour 710 (clone bg1fpj, 107669, Biolegend), PDL1-PE (clone 29E.2A3, 329705, Biolegend), CX3CR1-AF700 (clone SA011D11, cat# 149036, Biolegend), CD206-BV785 (clone MR5D3, cat# 565250, eBiosciences), Arg1-PE-Cy7 (clone A1exF5, cat# 25-3697-82, Thermo Scientific), Alox15 (cat# ab244205, Abcam), CD3-AF700 (clone 17A2, cat# 100216, Biolegend), CD4-V500 (clone RM4-5, cat# 100551, Biolegend), CD8-PE-CF594 (clone 53-6.7, cat# 562283, BD Biosciences). For intranuclear staining, the cells were treated with fixation/permeabilization buffer (eBioscience, 00-5123 and 00-5223) for 1h followed by intracellular staining. For intracellular staining the Cytofix/Cytoperm fixation/permeabilization kit (cat # 554714, BD Biosciences) was used. For Alox15 staining, the cells were incubated with AF488 conjugated donkey antirabbit antibody (cat#A32790, Thermoscientific) for 30 min at 4 C°. T cell subsets were identified with following antibodies: Tbet-PE (clone 4B10, cat# 644810, Biolegend), Rorγt-BV421 (clone Q31-378, cat# 562894, BD Biosciences), Gata3- Percp5.5 (clone 16E10A23, cat# 653812, Biolegend), and FOXP3-Pe-Cy7 (clone FJK-16s, cat# 15380970, eBiosciences). Cells were acquired using Aurora (Cytek) and analyzed using Flowjo software 10.10.0. For analysis of phagocytosis, pHrodo-labelled myelin or pHrodo labelled zymosan particles (Invitrogen, P35364) were used.

### Cell sorting

For sorting immune cells, the Neural Tissue Dissociation kit T (Miltneyi Biotec,130-093- 231) was used to generate brain/spinal cord homogenates following the manufacturer’s protocol. Subsequently, the homogenates were treated with percoll for myelin removal as described above. The cell suspensions from 4 mice per condition were pooled and then stained with PE anti-mouse CD45 (clone 30-F1, 1:200) and Live/Dead Fixable Near-IR Cell Stain Kit (Invitrogen, L34976, 1.500). CD45^+^ cells were sorted and collected using a SONY SH800 (SONY), BD Influx or FACS ARIAIII (BD biosciences) cell sorters. For May-Grünwald/Giemsa staining, CD45^+^ cells gated on the HG cell population were spun onto microscope glass slides using a cytospin and stained according to the protocol. For lipidomic analysis, the HG cells were sorted as SSC^high^CD45^+^CD11b^+^Ly6G- F4/80^lo^MHCII^lo^ using the Aurora sorter (Cytek). Control cells were sorted as SSC^lo^CD45^+^CD11b^+^Ly6G-Ly6C^-^.

### Lipid analysis

Sample Preparation: A specific sample amount was spiked with a mixture of antioxidants and internal standards consisting of 14,15-DHET-d11, 15-HETE-d8, 20-HETE-d6, 8,9- EET-d11, 9,10-DiHOME-d4, 12,13-EpOME-d4, 13-HODE-d4, LTB4-d4, with 500 pg each (Cayman Chemical, Ann Arbor, USA). Methanol was added. An alkaline hydrolysis was performed using sodium hydroxide at 60° C for 30 min. After centrifugation and pH adjustment, the obtained supernatant was added to Bond Elute Certify II columns (Agilent Technologies, Santa Clara, USA) for Solid Phase Extraction. The eluate was evaporated on a heating block at 40° C under a stream of nitrogen to obtain a solid residue. The residues were dissolved in 100 µL methanol/water.

LC/ESI-MS/MS: The residues were analyzed using an Agilent 1290 HPLC system with binary pump, multisampler and column thermostat with a Zorbax Eclipse plus C-18, 2.1 x 150 mm, 1.8 µm column using a gradient solvent system of aqueous acetic acid (0.05 %) and acetonitrile/methanol 50:50 (v/v). The flow rate was set at 0.3 mL/min, the injection volume was 20 µL. The HPLC was coupled with an Agilent 6495 Triplequad mass spectrometer (Agilent Technologies, Santa Clara, USA) with electrospray ionisation source. Analysis was performed with Multiple Reaction Monitoring in negative mode, with at least two mass transitions for each compound. Quantification was performed using calibration curves from authentic standards and their deuterated analogues.

### Phagocytosis assay *in vitro* and *ex vivo*

Primary microglia were cultured in 12-well plates with a density of 0.3 × 10^6^ per well as described above. The following day the medium was removed, and AEC-conditioned media (CM) was added for three days (see primary cultures). The control wells received regular microglia medium. Subsequently, the wells were rinsed with PBS and were exposed to pHrodo labelled zymosan bioparticles (Invitrogen, P35364) for 20min. The cells were washed with PBS, trypsinized with trypsin-EDTA 0.05% (Life Technologies, 25300-54) and subsequently stained according to procedure for flow cytometry.

To assess phagocytosis *ex vivo*, myeloid cells were isolated from PBS- or AEC-treated EAE brains at the disease peak as described above. The cells were resuspended in DMEM- F12 medium containing pHrodo (Life technologies, 36600) labelled myelin debri in the absence of FBS and incubated for 60min. Subsequently the cells were stained according to the procedure for flow cytometry and phagocytosis was assessed using Cytek Aurora.

### T cell recall stimulation

2D2 mice were injected with 25µg MOG peptide emulsified in Complete Freund’s Adjuvant as described above. On day 10, the mice were decapitated after CO_2_ inhalation. Mesenteric lymph nodes were collected, mechanically dissociated and pelleted using 350g centrifugation for 5 min at 4°C. The cells were resuspended in RPMI medium supplemented with 10% FBS, 2mM l-glutamine, 100U/mL penicillin, and 100μg/mL streptomycin. AECs were added to the lymph node cells at ratios of 1:2 or 1:4 AEC:Lymph node cells. The cells were stimulated with MOG peptide (Anaspec, 60130-5) at a concentration of 20µg/mL and incubated for 72 hours at 37°C and 5% CO_2_. The cells were collected, washed with PBS and stained with Live/Dead marker in the presence of Fc block (Invitrogen, 14-0161-83, 1:200). For surface staining the following mouse antibodies were used: CD3-PE/Dazzle-594 (Biolegend, 317345, [Clone OKT3],1:100), CD4-FITC (Biolegend, 357406, [clone A161A1], 1:100), CD8-V500 (BD, 560776, [Clone 53-6.7],1:100), CD25-PE and (Biolegend, 102010, [clone PC61], 1:100) for 30 min. The cells were washed with FACS buffer (PBS, 1% FBS, 2mM EDTA) and fixed and permeabilized according to the Foxp3/transcription factor staining buffer set (eBioscience, 00-5523-00) for intracellular staining. The cells were stained with anti-mouse antibodies including FOXP3-AF700 (eBiosciences, 25-5773-82, clone FJK-16s, 1:100), IFNγ-APC (Biolegend, 554413, clone XMG1.2, 1:100), TNFα-PE-Cy7 (eBiosciences, 25-5773-82, clone TN3- 19.12), IL-17A-Percp-Cy5.5 (eBiosciences, 45-7177-82, clone eBio17B7) and IL10- BV421 (Biolegend, 505021, clone JES5-16E3) for 1 hour in 4 °C. Subsequently, the cells were washed and acquired using an Aurora (Cytek).

### Reverse transcription polymerase chain reaction (RT-PCR)

Total RNA of cells or CNS tissue was obtained using the RNeasy mini kit (Qiagen, 74106) with on-column DNase I digestion using (Qiagen, 79254) according to the manufactoŕs instructions. RNA was reverse transcribed to cDNA using iScript kit (BioRad Laboratories, 1708891). Amplifications were conducted in a 384 well plate using SYBR green (BioRad Laboratories, 1708886) and run in BioRad CFX384 Touch Real-Time PCR Detection System. Primer specificity was determined by melt curve analysis of each reaction indicating a single peak, and annealing was obtained at approximately 60°C. The following primers were used: *Hrpt*: 5’(ACAGCCCCAAAATGGTTAAGG); 3’ (TCTGGGGACGCAGCAACTGAC), *Gapdh*: 5’(TGAAGCAGGCATCTGAGGG); 3’ (CGAAGGTGGAAGAGTGGGAG). *Nos2:* 5’(GTTCTCAGCCCAACAATACAAGA); 3’ (GTGGACGGGTCGATGTCAC). *Il1b*: 5’(GCAACTGTTCCTGAACTCAACT); 3’ (ATCTTTTGGGGTCCGTCAACT). *Il6*: 5’(TAGTCCTTCCTACCCCAATTTCC); 3’ (TTGGTCCTTAGCCACTCCTTC). *Tnfa*: 5’(CTGTAGCCCACGTCGTAGC); 3’ (TTGAGATCCATGCCGTTG). *Arg1*: 5’(CTCCAAGCCAAAGTCCTTAGAG); 3’ (AGGAGCTGTCATTAGGGACATC).

### Cytometric bead array

Cytokine release to the supernatants was measured using BD™ Cytometric Bead Array (CBA) according to the instructions provided by the manufacturer. The beads were detected using a BD LSRII flow cytometer. The following cytokines were analyzed: Mouse IL-6 Flex Set (BD, 558301), Mouse TNF Flex Set (BD, 558299), and Mouse IL-10 Flex Set (BD, 558300).

### Immunofluorescent staining

For immunofluorescence, cells were plated in 96-well plates at density of 1x10^4^ cells per well. Following treatment (see above) the cells were fixed in 100μL 4% PFA for 30min at room temperature. After 3 consecutive washes in PBS, the cells were incubated with blocking buffer (PBS containing 0.3% Triton X-100 and 5% normal goat or donkey serum) for 30 min at room temperature. After removal of the blocking solution, the cells were incubated with primary antibodies diluted in blocking buffer overnight at +4°C. The following antibodies were used: Iba1 (Wako, 019-19741, 1:400), Ly6C (Abcam, ab54223 1:100), CD55 (Invitrogen, PA5-88462, 1:100), HLAG (EXBIO, 1A-292-C100, 1:100), K8/K18 (Progen Biotech, 10502, 1:50), Arg1 (Abcam, ab60176 1:100), Alox15 (Abcam, ab244205, 1:125), MHCII (Invitrogen, 14-5321-82, 1:200), P2RY12 (Generated by Dr. O.

Butovsky, 1:400). After washing 3 times with PBS, the cells were incubated with the corresponding secondary antibodies for 2 hours at room temperature. The cells were washed with PBS, and counterstained with Hoechst for 10min (Thermo Scientific, 62249, 1:10000). Perfused brains/spinal cords or whole skulls were fixed in 4% paraformaldehyde. The whole skulls were decalcified in 120mM EDTA for 1 week at 37°C. The tissues were dehydrated overnight in 20% sucrose, embedded in OCT, and frozen on dry ice. Using a cryostat (Invitrogen, CM1850), 14μm thin sections were prepared. Staining was performed as indicated above. Confocal images were acquired with a Zeiss LSM880 microscope and analyzed using ImageJ default plugins. The morphological analyses of Arg1^+^ macrophages was performed using Imaris 9.8 with the Surface function, followed by segmentation. The Sphericity and Volume of each segmented Arg1^+^ and Arg1^-^ macrophages were generated using the built-in statistics feature of Imaris 9.8.

### Single cell RNAseq

The transcriptomics libraries were prepared using Chromium Next GEM Single Cell 3 Reagent Kits v3.1 (Dual Index) (10X genomics) using the protocol provided by the manufacturer (CG000315 Rev C). Concentration of the libraries was estimated using Qubit™ dsDNA Quantification High Sensitivity Assay Kit (Invitrogen) and KAPA Library Quantification Kit (Roche). Quality control on the libraries was performed using Bioanalyzer High Sensitivity DNA Kit (Agilent). Samples were sequenced on NextSeq2000 (NextSeq 1000/2000 Control Software 1.4.1.39716/RTA 3.9.25) with a 28nt(Read1)-10nt(Index1)-10nt(Index2)-90nt(Read2) setup using ’P2’ flowcell. The HG samples were sequenced on NovaSeqXPlus (control-software 1.1.0.18335) with a 151nt(Read1)-19nt(Index1)-12nt(Index2)-151nt(Read2) setup using ’10B’ mode flowcell. The Bcl to FastQ conversion was performed using bcl2fastq_v2.20.0.422 from the CASAVA software suite. The quality scale used is Sanger/phred33/Illumina 1.8+.

### Single cell RNA-seq data processing

Sequenced samples were processed via the Cell Ranger pipeline (version 7.0.1, 10x Genomics). Reads were aligned to GRCm39 mouse reference genome.

Downstream analysis was performed in RStudio (R version 4.3.2) using package Seurat (versions 5.0.1 and 5.1.0). Cells expressing fewer than 200 genes or greater than 20% mitochondrial reads were excluded from the analysis in both brain datasets (CD45^+^ high granular (HG), and CD45^+^ cells). In the spinal cord dataset, cells expressing fewer than 100 genes were excluded while the 20% mitochondrial read threshold was kept. Doublets were removed using DoubletFinder.

After quality control (QC), the counts were normalized and scaled regressing out the mitochondrial genes and total counts per cell. FindVariableFeatures and SelectIntegrationFeatures functions from Seurat package were used to select variable genes. Top 50 principal components were calculated and used for data integration to diminish the batch effect between different samples and datasets. Harmony was used to integrate two samples within the same dataset (e.g. brain CD45^+^ dataset) as well as combining two datasets (e.g. brain CD45^+^ dataset and CD45^+^HG dataset). Then the UMAP and Louvain clustering were performed. Wilcox test embedded in FindAllMarkers function was used to identify differentially expressed genes (DEGs) for each cluster with a log2 fold change larger than 0.2, and an adjusted p-value smaller than 0.05. Top5 genes were selected in an unsupervised manner by ranking genes using percentage difference and adjusted p-value. DEGs were used to annotate cell types and isolated cell types were re-clustered based on the DEGs to identify new subtypes. The Seurat build-in function AddModuleScore was used to assign scores to individual clusters based on gene lists from indicated publications. The gene list for Phagocytosis score was taken from Gene Ontology term ‘positive regulation of phagocytosis’ (GO0050766).

The python software SCENIC^13, 14^ was used to predict regulon activity (performed in conda environment with python 3.10.14). Gene regulatory network was inferred with the GRNBoost2 algorithm in the combined datasets of Brain. Then, candidate regulons were predicted by cisTarget. AUCell was employed for predicting the cellular enrichment of predicted regulons. Top regulons were identified by the build-in function regulon_specificity_scores. The activity of each regulon was binarized as ‘on’ or ‘off’ state for visualization.

Gene set enrichment analysis (GSEA) was performed on DEGs for each subtype via the function GSEA from R package ClusterProfiler (version 4.12.6, built on fgsea). Fold change was used to rank GSEA list. Dotplots were generated by dotplot function from R package enrichplot (version 1.24.4).

Cellular interactions were imputed by R package CellChat (version 2.1.2) after combining the annotated cell subtypes from Brain CD45^+^ dataset and brain HG datasets and then splitted by condition (PBS vs AEC). CellChat built-in function netVisual_heatmap was used to generate the heatmap comparing interaction number and interaction strength between the two conditions. To compare pathway-specific cellular interaction between the two conditions, the function netVisual_chord_gene or netVisual_heatmap was used to generate chord graph, and heatmap respectively.

R package slingshot (monocle3 for verification) was used to infer the pseudotime of monocyte, macrophage and HG3 activation path. Trajectories were imputed based on 3D UMAP after integrating the following clusters using harmony: HG3, Mac1, Mac2, Mon1, Mon2, Mon3, Mon4, and Mon5. Lineage-associated genes were calculated by R package Tradeseq (version 1.4) based on lineages inferred by Slingshot.

### Bulk RNAseq and data analysis

Wildtype mice received intracisternal injection of AECs or PBS as described above. Brains were collected after cardiac perfusion with PBS 6 hours, 1 day and 3 days post AEC- transfer. Control mice received PBS. The brains were stored in RNAprotect tissue reagent overnight (2 mL/brain, Cat #:76104, Qiagen). The leptomeningeal layer was scrapped and collected together with choroid plexus in 1.5 mL tube. RNA was extracted from all samples using the RNeasy Mini Kit (QIAGEN) as described above, and RNA integrity was assessed using an Agilent 2100 Bioanalyzer (Agilent Technologies). mRNA-seq library preparation and sequencing were performed on Illumina platforms (Novogene Biotechnology Co., Nanjing, China) with 150-base pair paired-end reads. The raw RNA sequence files were processed to extract FASTQ data, which underwent quality control (QC) checks using FastQC to filter low-quality reads. Reads with adaptor contamination, more than 10% uncertain nucleotide content, or 50% low quality nucleotides were filtered out. After the QC, the reads were aligned to the mouse reference genome (GRCm38 mm10 assembly) using HISAT2 (version 2.0.5). Gens with count<1 were removed resulting in 24688 genes for downstream analysis. R package DESseq2 (version 1.44.0) was used to identify DEGs. Functions ggvenn (version 0.1.15) and Enhanced Volcano (version 1.22.0) in R package was used for generating Venn and volcano plots. Using IfcShrink function the DEGs were ranked for subsequent GSEA analysis as described above.

### Statistical data analysis

GraphPad Prism 9.5.1 software was used to perform statistical analysis and generate graphs. Comparison between two groups was performed with Student’s two-tailed unpaired *t* test or Mann Whitney test. For multiple comparisons one-way or two-way ANOVA with subsequent Tukey’s multiple comparison test was performed. *P* < 0.05 was considered statistically significant.

## Acknowledgements

We thank the following core facilities for their assistance: CMM FACS core facility (Annika van Vollenhoven), Biomedicum Flow Cytometry (BFC, Alda Saldan), Biomedicum imaging core (BIC), CMM fFMI, CMM single cell facility core. We acknowledge support from the National Genomics Infrastructure in Stockholm funded by Science for Life Laboratory, the Knut and Alice Wallenberg Foundation and the Swedish Research Council, and NAISS/Uppsala Multidisciplinary Center for Advanced Computational Science for assistance with massively parallel sequencing and access to the UPPMAX computational infrastructure. The data handling was enabled by resources in project [NAISS 2024/23-284 and NAISS 2023/22-1282] provided by the National Academic Infrastructure for Supercomputing in Sweden (NAISS) at UPPMAX, funded by the Swedish Research Council through grant agreement no. 2022-06725.

## Author contributions

HS and RAH conceptualized the study with input from RG on AECs. HS designed all the experiments, performed most of the experiments and analyzed the data. YG performed RNAseq data processing and analysis, assisted with experiments. SB (confocal microscopy, AEC isolation); KZ (cell sorting, scoring, Imaris analysis, assistance with experiments); VJ (single-cell library preparation); GV (EAE scoring, tissue processing); YC (preparation of human MDMs); JHM (recall experiment, scoring, tissue preparation); IBC (cryosectioning, tissue preparation); UR and OGT (preparation of human T cells); JL (confocal microscopy and tissue preparation); VS and PTA (in vitro phagocytosis assay, recall experiment), AOGC (MOG preparation, input on T cells and recall experiment). RG (AEC cultures, AEC isolation). HS wrote the manuscript, and all the authors contributed to evaluation of the results and discussions in writing of the manuscript.

