## Supplementary Figures for "Human placental stem cells induce a novel multiple myeloid cell-driven immunosuppressive program that ameliorates proinflammatory CNS pathology"

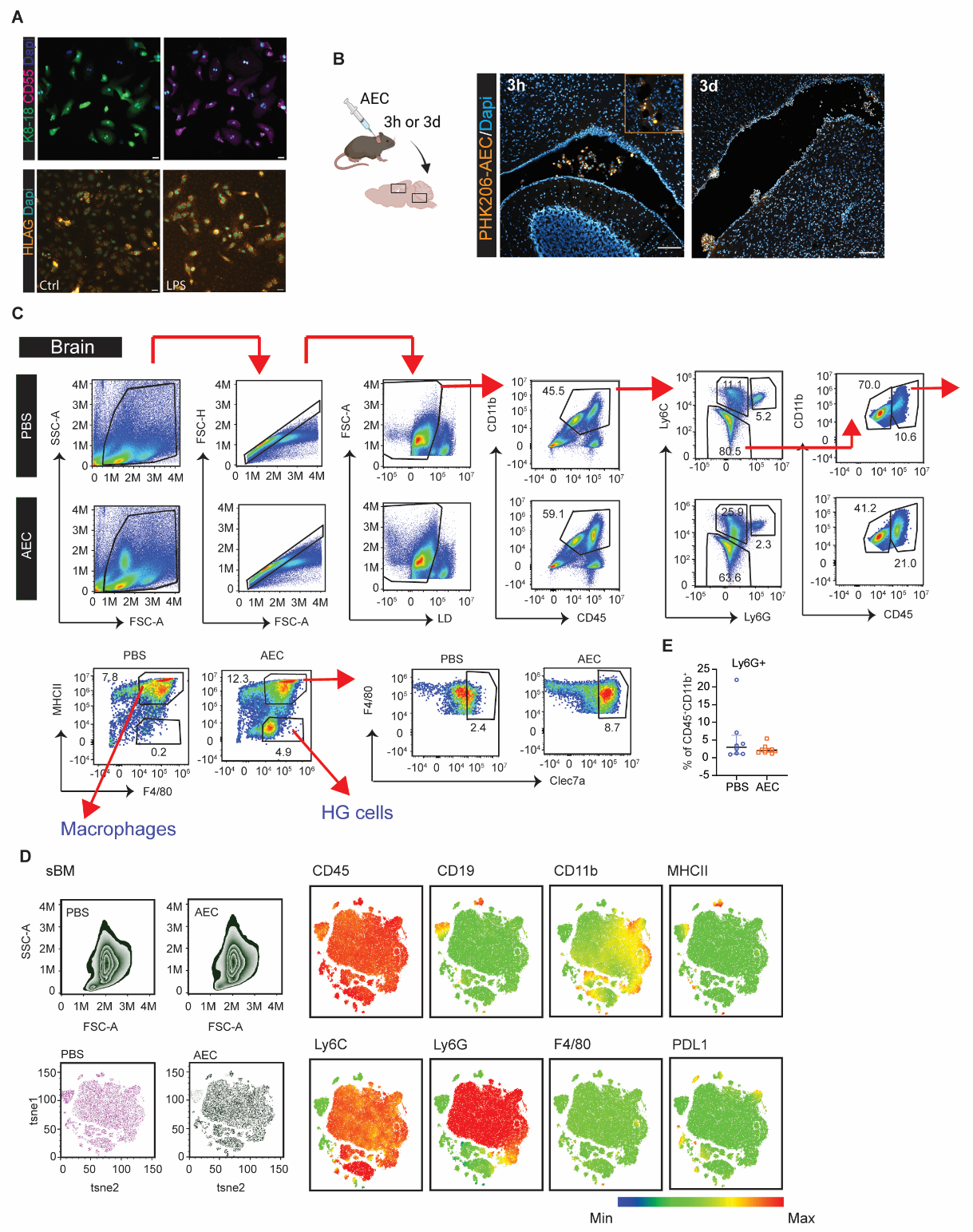

**Figure S1. AEC characteristics and their presence in the brain post-AEC injection.** **A** Representative immunofluorescence image of CD55, K8/18 and HLA-G in AECs. HLA-G expression is typically reduced *in vitro* but is inducible by LPS. Scale bar=20µm. **B** Experimental design of AEC tracking. Representative immunofluorescence image of PHK206-labelled AECs in the brain 3 hours and 3 days post -AEC injection. **C** Gating strategy of the myeloid cells in EAE brains of PBS- and AEC-treated mice. The average proportion of each population within total myeloid cells (CD45^+^CD11b^+^) is indicated in each plot. **D** tSNE plots of skull bone marrow indicating the expression of myeloid markers and CD19 in PBS- or AEC-treated EAE mice. **E** Quantification of Ly6G^+^ cells in the brains of PBS- and AEC-treated EAE mice. n=8 (PBS), n=9 (AEC). Data is presented as mean±SD; statistical analysis: unpaired Student’s t-test. **p* < 0.05; ***p* < 0.01

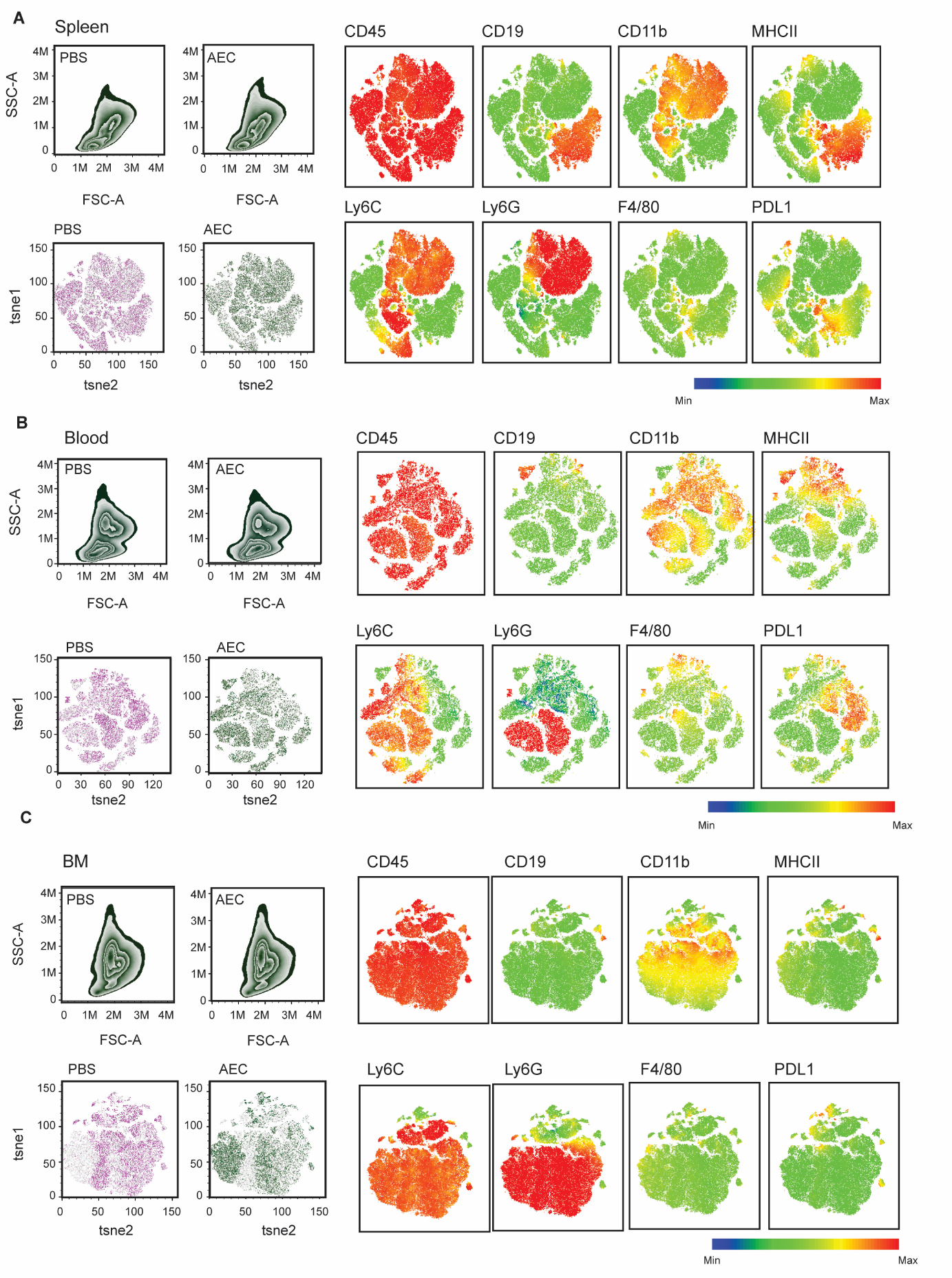
**Figure. S2** **Intracisternal AEC treatment does not affect peripheral immune cells.** tSNE plots of **A** spleen **B** blood **C** bone marrow indicating the expression of myeloid markers and CD19 in PBS- or AEC-treated EAE mice.

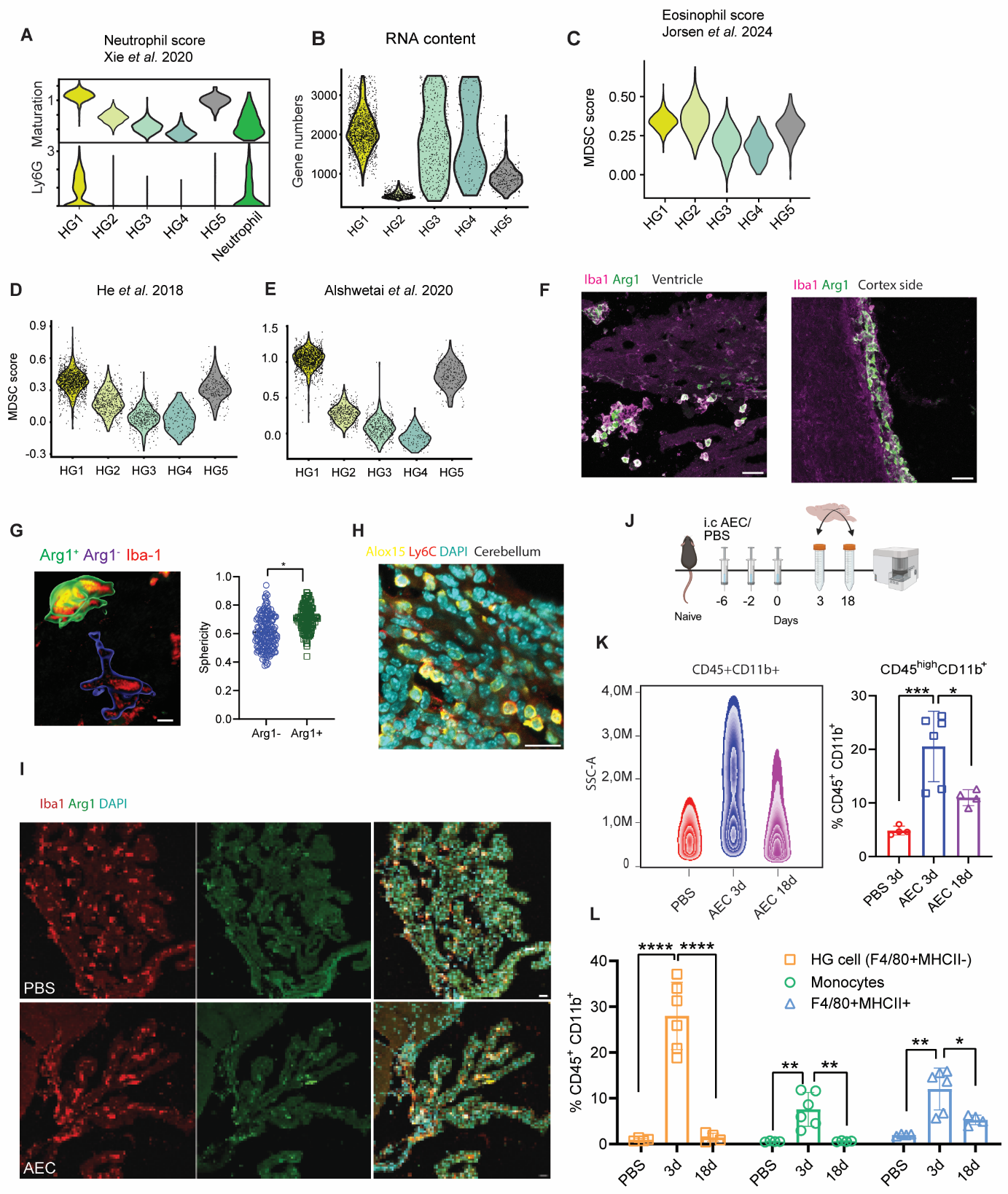
**Figure S3. The impact of AEC treatment on myeloid subsets in the brain.** **A** Violin plot indicating neutrophil score in clusters of high granular cells. **B** Violin plot indicating low RNA content in HG2 as indicated by low gene number. **C** Violin plot indicating eosinophil score in clusters of high granular cells. **D** Violin plot indicating a comparison of HG clusters with MDSC score in developmental and **E** breast cancer context. **F** Representative image of Arg1 immunofluorescence in the brain borders. Scale bar=40µm. **G** Morphological analysis of macrophages expressing Arg1 using Imaris. A total of 171 Arg1^-^ and 144 Arg1^+^ macrophages were analyzed in n=4 AEC choroid plexus. Data are presented as mean ± SD; statistical analysis: unpaired Student’s t-test comparing n=4 Arg1^+^ vs n=4 Arg1^-^ condition. **H** Representative image of ALOX15 immunofluorescence in parenchyma close to 4^th^ ventricle. Scale bar=40µm. **I** Representative expression of ARG1 and Iba1 in choroid plexus at the end of EAE. Scale bar= 40µm. **J** Experimental design of AEC treatment. **K** FACS plot indicating CD45^+^CD11b^+^ myeloid cells and the quantification of CD45^high^CD11b^+^ myeloid cells 3 and 18 days after last AEC injection. **L** Quantification of myeloid subsets within CD45^high^CD11b^+^ compartment. Data is presented as mean±SD; statistical analysis: One-Way ANOVA followed by Tukey’s t-test. n=4 PBS, n=6 AEC 3d, n=4 AEC 18d.

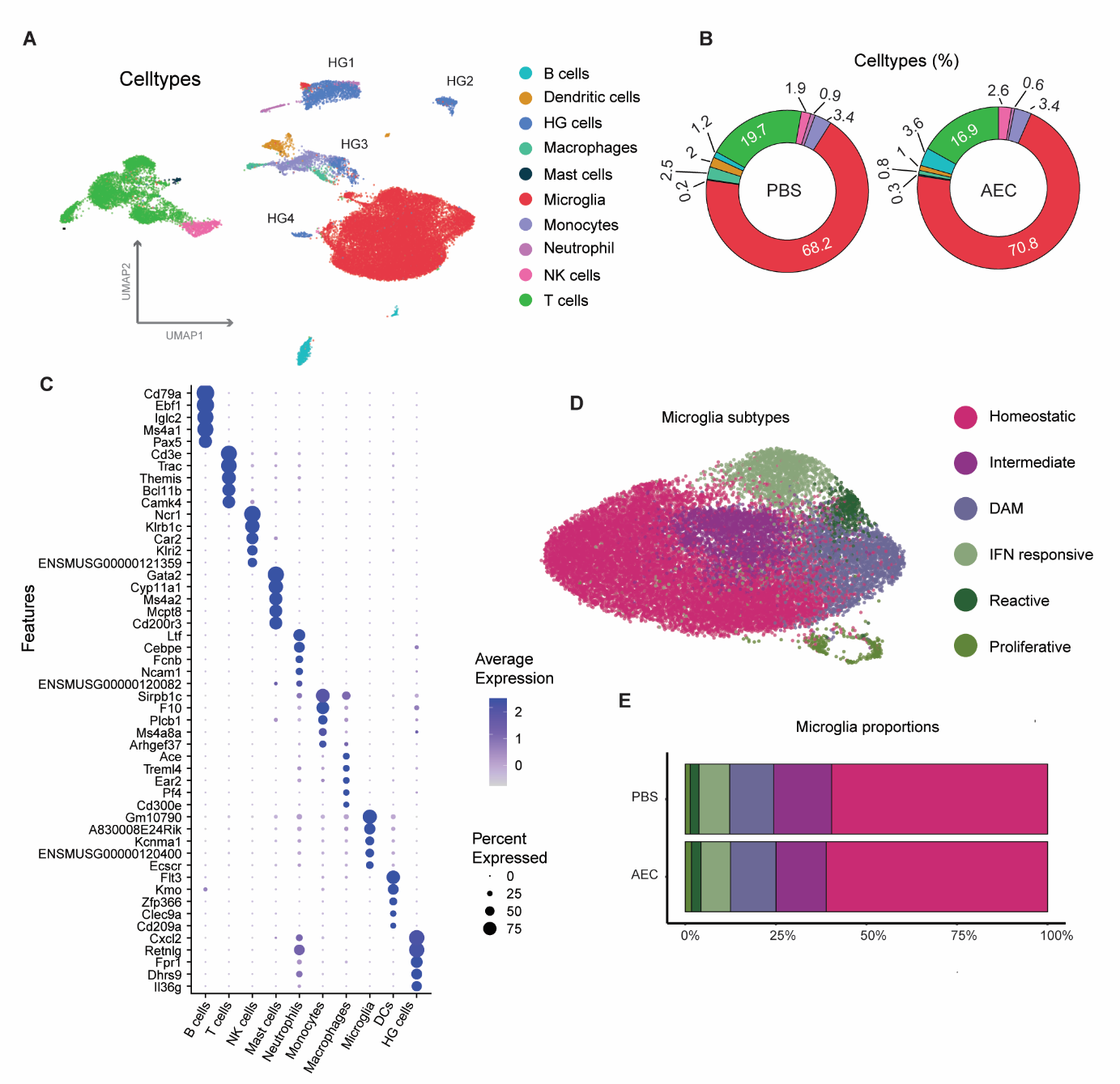

**Figure S4.** **AEC-induced myeloid cells in the brain are transient**. **A** UMAP of 30677 CD45^+^ immune cells and 2858 high granular cells (blue) projected together. **B** The proportion of CD45^+^ immune cells in each condition. **C** Top5 hallmarks of each cluster. **D** UMAP of 21306 microglia cells and **E** the proportion of their subsets in AEC and PBS conditions.

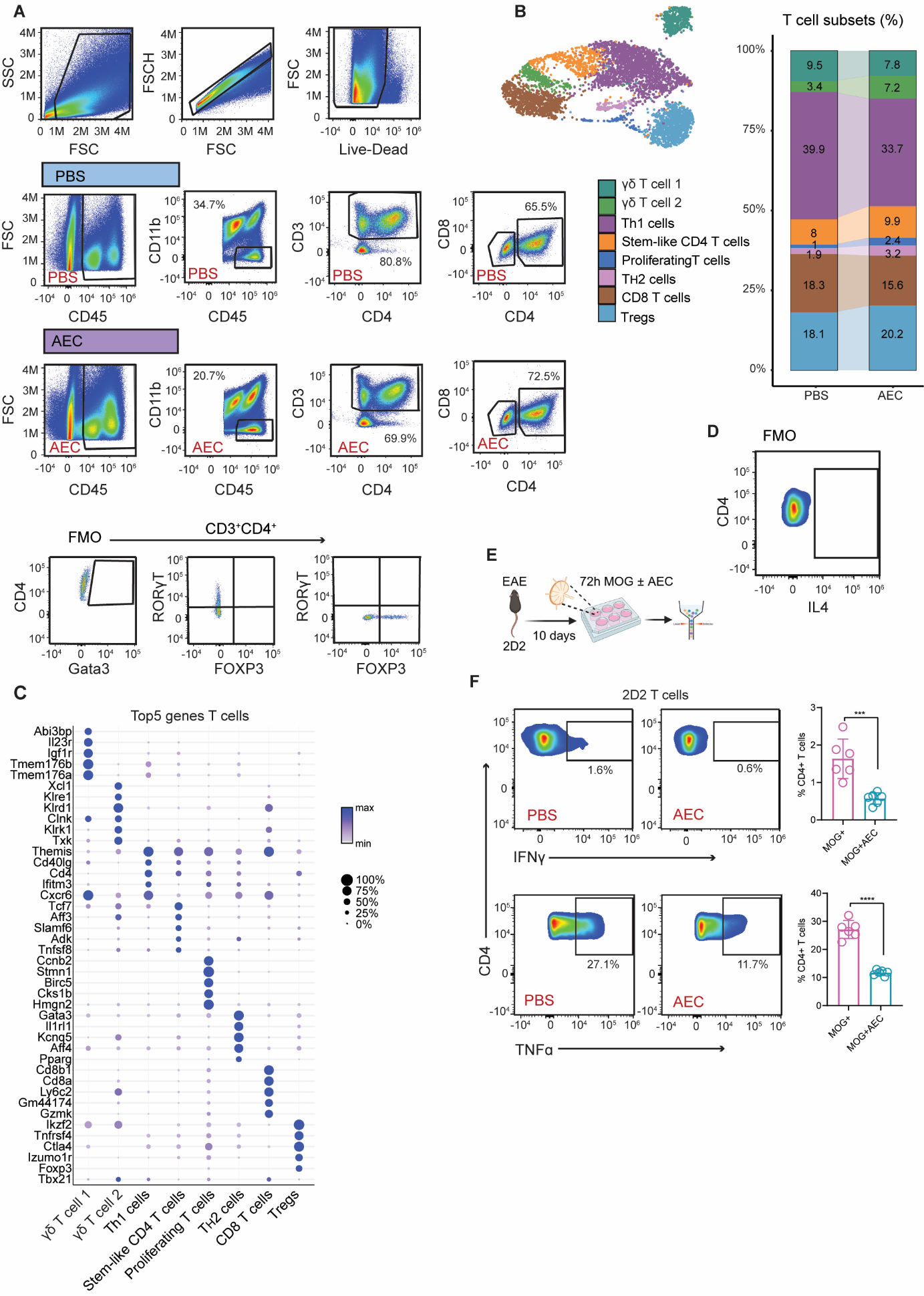

**Figure S5. The effects of AEC treatment on brain T cells. A** Gating strategy of T cells in EAE brains of PBS- and AEC-treated mice. Averages of each population are indicated in the graph. Fluorescence minus one (FMOs) are indicated in the lowest panel. **B** UMAP of 5619 T cells obtained from EAE brains and their proportion in each condition. **C** Top5 hallmarks of each T cell cluster. **D** FMO for IL4 in CD4 T cell subset. **E** Experimental design of Recall experiment. **F** FACS plot indicating cytokine expression in CD4^+^ T cells after treatment with PBS or AEC in lymph node extracts derived from MOG immunized 2D2 mice. Quantification is indicated to the right and the averages in the respective plot. Data are presented as mean ± SD; n=6 PBS, n=6 AEC. statistical analysis: unpaired Student’s t-test. **p* < 0.05; ***p* < 0.01; ****p* < 0.001

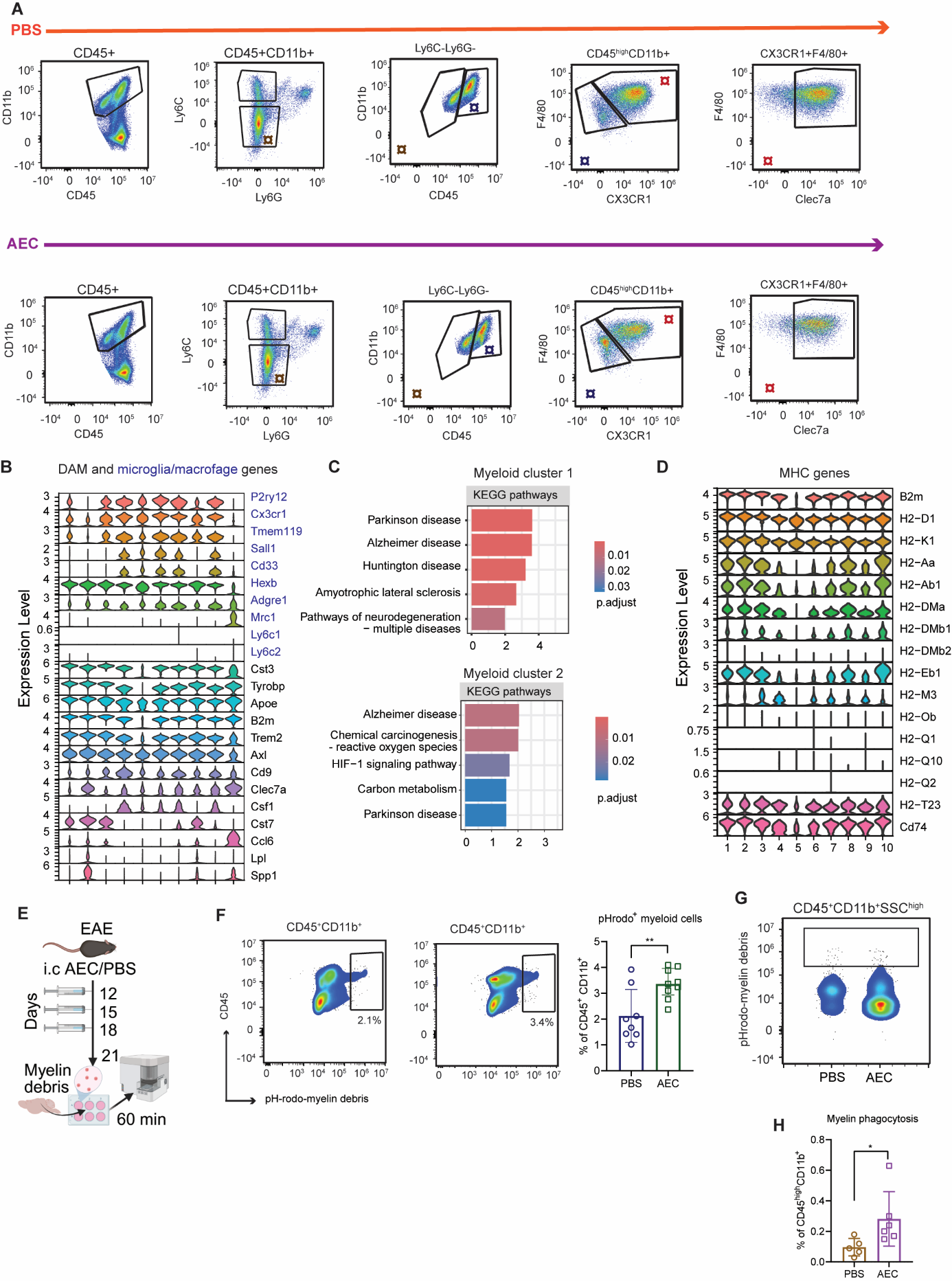

**Figure S6. The effects of AEC treatment on spinal cord. A** Gating strategy of the myeloid cells in EAE spinal cord of PBS- and AEC-treated mice after excluding single cells and dead cells. **¤** indicates gated populations. **B** Violin plots indicating DAM genes and microglia/macrophage genes within the myeloid subsets in the spinal cord. **C** KEGG pathways for Cluster 1 and Cluster 2 in the spinal cord. **D** Violin plots indicating MHC genes within the myeloid subsets in the spinal cord. **E** Experimental design of *ex vivo* phagocytosis experiment. **E** FACS plot indicating myelin uptake in CD45^+^CD11b^+^ myeloid cells in the brain with averages indicated in the plot. Data is presented as mean ± SD; n=8 PBS and n=9 AEC. statistical analysis: unpaired Student’s t-test. **F** FACS plot indicating high granular CD45^+^CD11b^+^ myeloid cells and their minimal involvement in myelin uptake. **H** Quantification of myelin uptake by macrophages in the spinal cord. Data is presented as mean ± SD; statistical analysis: unpaired Student’s t-test. n=5 PBS and n=6 AEC. **p* < 0.05; ***p* < 0.01; ****p* < 0.001
